# Plasticity in prefrontal cortex induced by coordinated nucleus reuniens and hippocampal synaptic transmission

**DOI:** 10.1101/2020.07.11.197798

**Authors:** Paul J Banks, E Clea Warburton, Zafar I Bashir

## Abstract

The nucleus reuniens of the thalamus (NRe) is reciprocally connected to a range of higher order cortices including hippocampus (HPC) and medial prefrontal cortex (mPFC). The physiological function of NRe is well predicted by requirement for interactions between mPFC and HPC, including associative recognition memory, spatial navigation and working memory. Although anatomical and electrophysiological evidence suggests NRe makes excitatory synapses in mPFC there is little data on the physiological properties of these projections, or whether NRe and HPC target overlapping cell populations and, if so, how they interact. We demonstrate in *ex vivo* mPFC slices that NRe and HPC afferent inputs converge onto more than two-thirds of layer 5 pyramidal neurons, show that NRe, but not HPC, undergoes marked short-term plasticity at theta, and that HPC, but not NRe, afferents are subject to neuromodulation by acetylcholine acting via muscarinic receptor M2. Finally, we demonstrate that pairing HPC followed by NRe (but not pairing NRe followed by HPC) at theta frequency induces associative, NMDA receptor dependent synaptic plasticity in both inputs to mPFC. These data provide vital physiological phenotypes of the synapses of this circuit and provide a novel mechanism for HPC-NRe-mPFC encoding.

---

The medial prefrontal cortex (mPFC) is vital for performance of many higher order cognitive functions including decision making, attention, and mnemonic processing. Evidence has emerged that communication between mPFC and the hippocampus (HPC), which results in synchronous oscillatory activity and cell firing (Jones & Wilson 2005, Siapas et al 2005), is required for performance of some of these behaviours including during the encoding phase of spatial working memory (Spellman et al 2015) and associative recognition memory (Barker et al 2017).

More recently, regions of the ventral midline thalamus centred upon the nucleus reuniens (NRe) have been shown to be involved in many cognitive processes which require HPC-mPFC interactions, including encoding of associative recognition memory (Barker & Warburton 2018) and working memory (Hallock et al 2016). NRe has dense reciprocal connections to both HPC and mPFC and has been proposed to be a primary route of feedback from the mPFC to HPC (Cassel et al 2013). NRe relays trajectory information from mPFC to CA1 during spatial navigation (Ito et al 2015). NRe is also reciprocally connected to associative cortices including entorhinal and perirhinal cortex, which are themselves directly and indirectly connected to HPC and mPFC. Deactivation of NRe reduces oscillatory coherence and phase-locking between HPC and mPFC during spatial working memory (Hallock et al 2016), suggesting that NRe is situated ideally as a long-range coordinator of higher order cortical structures.

Moreover, manipulations of NRe produce complex delay-dependent effects on HPC-mPFC dependent memory, suggesting that NRe does not act as a simple relay but is important for encoding memory by coordination of long-range connections (Barker & Warburton 2018). Indeed, the behavioural effects of blocking protein synthesis within NRe suggest that plasticity processes may even encode and store memory within NRe itself (Barker & Warburton 2018).

Although evidence for the functional importance of NRe in cognitive processes is strong, very little is known about the nature of NRe afferents to mPFC, or how they may interact with those of the HPC (Di Prisco & Vertes 2006, Eleore et al 2011). It is unknown whether NRe and HPC synapse onto overlapping mPFC cell populations and whether these synapses have divergent receptor expression, temporal properties or are subject to differential neuromodulation and plasticity. Answering these questions is key to understanding how HPC and NRe input is assimilated in mPFC.

Here we use an optogenetic strategy to detail the physiological properties of NRe synapses to pyramidal neurons in *ex vivo* mPFC brain slices. We show that NRe and HPC synapses converge on to the majority of L5 pyramidal neurons and we directly compare synaptic physiology of the distinct inputs into mPFC. We demonstrate marked differences in short-term plasticity between NRe and HPC and explore their regulation by neuromodulators acetylcholine (ACh) and dopamine, both of which have prominent roles in mPFC physiology. Based on the connectivity of the HPC, NRe and mPFC a simple circuit would envisage HPC driving mPFC directly but also driving mPFC indirectly with a delay via NRe. Alternatively, NRe can drive mPFC directly but also drive mPFC indirectly with a delay via HPC. Since synaptic plasticity in mPFC is considered important for learning (Meunier et al 2017, Sabec et al 2018) but little is known about cooperativity of inputs to mPFC, we finally demonstrate that patterned HPC and NRe synaptic activity in which HPC leads NRe (but not in the opposite direction) can interact to induce associative, NMDA receptor-dependent long-term plasticity. This unidirectional cooperativity has important implications for encoding of information within mPFC.

## Materials and Methods

### Animals

All experiments were carried out in naïve male Lister Hooded rats (Envigo, UK) weighing 300-450g at the start of experiments. Animals were housed in groups of 2-4 under a 12 h/12 h light/dark cycle with lights on 20:00-08:00 and were given ad libitum access to food and water. Sacrifice for ex-vivo slices occurred 2-3 hours into the dark cycle. All animal procedures were conducted in accordance with the United Kingdom Animals Scientific Procedures Act (1986) and associated guidelines. All efforts were made to minimise suffering and number of animals used.

### Viral injections

Optogenetic transduction of neurons was achieved using AAV9-CaMKii-hChR2(E123T/T159C)-mCherry (Addgene 35512; 3.3 × 10^13^ genome copies/ml) obtained from University of Pennsylvania Vector Core or Addgene. The viral vector was chosen due to: 1) the AAV9 serotype having previously been described as having few deleterious effects on synaptic release properties (Jackman et al 2014); 2) suitability of the CaMKii promotor for transduction of excitatory neurons present in NRe (Bokor 2002); 3) the hChR2(E123T/T159C) “ChETA_TC_” channelrhodopsin variant having a combination of fast kinetics and large photocurrents suitable for activation of distal sites (Berndt et al 2011, Mattis et al 2011). Each rat was anaesthetised with isoflurane (4% induction, 2.5-3.5% maintenance) and secured in a stereotaxic frame with the incisor bar set 3.3mm below the interaural line. For NRe injections bilateral burr holes were made in the skull at the following coordinates with respect to bregma: anterior-posterior (AP) – 2.0 mm, mediolateral (ML) **±** 1.4 mm. Virus was front loaded into a 33-gauge 12° bevelled needle (Esslab) attached to a 5 µl Hamilton syringe which was mounted at a 10° angle in the mediolateral plane to avoid the sinus, with the eyelet of the needle facing medially. The needle was lowered 7.5mm below the surface of the skull measured from the burr hole and 100 nl of virus was delivered via each burr hole at a rate of 200 nl.min^-1^, with the needle left in situ for 10 minutes after each injection. NRe viral injections transduced neurons approximately ± 1.0 mm in the anteroposterior axis. Injections produced strong mCherry expression in nucleus reuniens, rhomboid nucleus, and xiphoid with sparse expression in adjacent ventral reuniens (also referred to as the perireuniens), paraxiphoid and submedius thalamic nuclei (Fig 1A). Of those nuclei, only the reuniens, ventral reuniens and rhomboid nuclei project to prelimbic cortex (Alcaraz et al 2015, Hoover & Vertes 2007, Salay et al 2018); for simplicity we shall subsequently refer to these collectively as NRe. For HPC viral injections, coordinates were AP −6.3 mm, ML **±** 5.5 mm, dorsoventral −5.8 mm, measured from bregma. Hippocampal viral injection volume was 500 nl per hemisphere.

**Figure 1.**
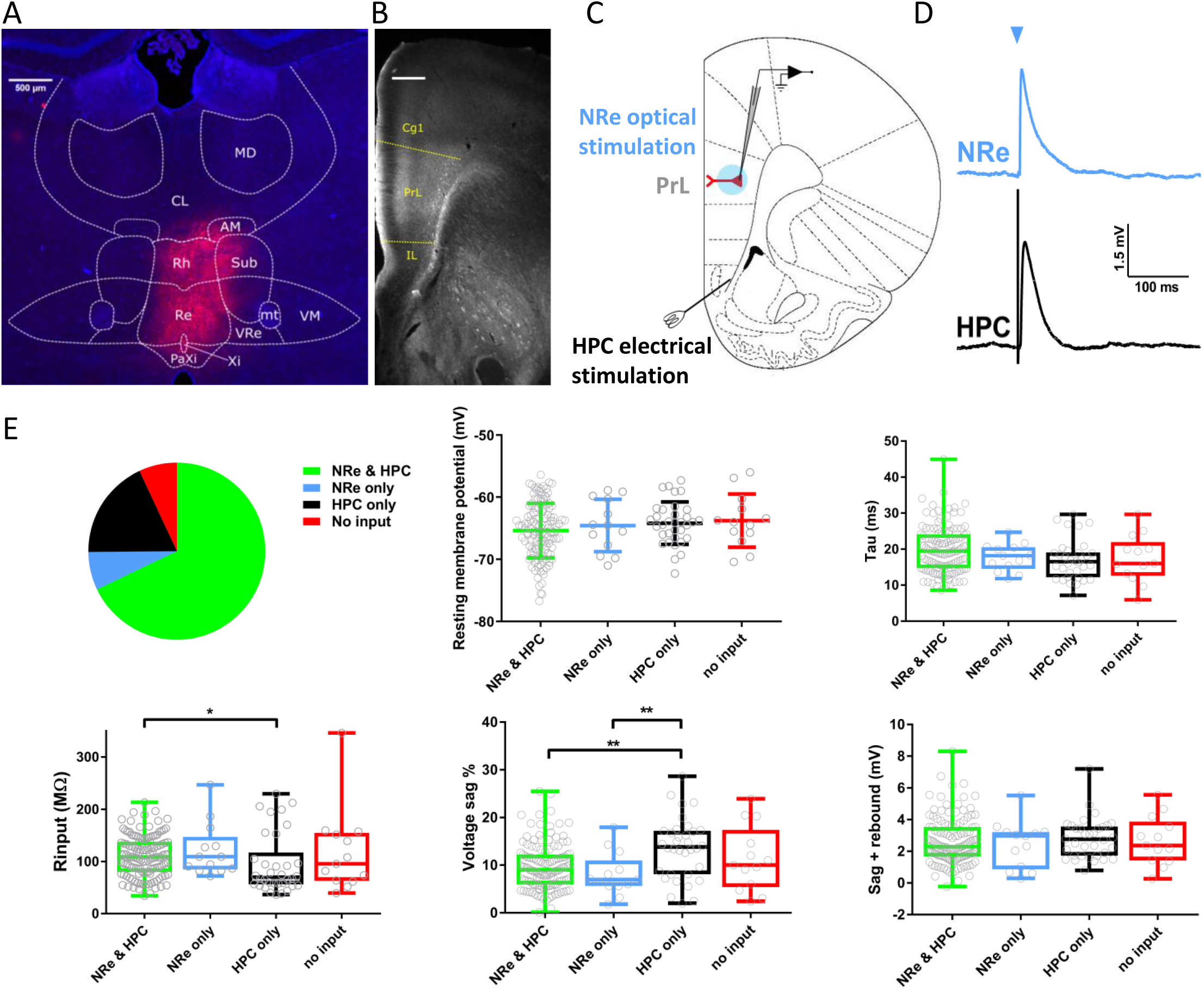
Electrophysiological characterisation of L5 pyramidal cells receiving optogenetically activated nucleus reuniens synapses. **A -** Representative widefield-fluorescence image showing neuronal transduction following injection of AAV9:CaMKii:hChR2(E123T/T159C):mCherry (red) into nucleus reuniens (Re) and DAPI (blue). VRe, ventral reuniens; Rh, rhomboid; Xi, xiphoid; PaXi, paraxiphoid; CM, central medial; AM, anteromedial; VM, ventromedial; MD, mediodorsal; Sub, submedius thalamic nuclei; mt, mammillothalamic tract. **B -** Monochrome image of mCherry positive fibres in PFC following AAV injection into nucleus reuniens. Dotted lines denote the boundaries of prelimbic cortex. mCherry signal is amplified with anti-mCherry antibody. Cg1 = cingulate cortex, IL = infralimbic cortex, PrL = prelimbic cortex. Scale bar = 500 µm. **C -** Schematic of acute mPFC slice with whole-cell recording from layer 5 pyramidal neuron in PrL, light activation of soma and proximal dendrites via microscope objective (blue) and stimulation of hippocampal fibre bundle using conventional stimulating electrode. **D -** Representative NRe (blue) and HPC (black) EPSPs. Blue arrow denotes light activation. **E –** Proportion of cells receiving different permutations of NRe and HPC inputs, 187 cells from 65 animals. Passive membrane properties measured from −100 pA current injection split by synaptic input. Resting membrane potential plotted as mean ± SD, one-way ANOVA F_(3,183)_ = 1.2, p = 0.32. Other parameters one or more column failed Shapiro-Wilk test for normality, box plots show median and interquartile range, whiskers max and min data points. Kruskal-Wallis test p values: Tau = 0.074, Rinput = 0.031, Sag % = 0.0036, sag + rebound = 0.84. */** = p <0.05/0.01 Dunn’s multiple comparisons post-hoc.

### Acute slice preparation

After a minimum of 10 days following viral injection, animals were anaesthetised with 4% isoflurane and decapitated. Brains were rapidly removed and placed into ice-cold sucrose solution (in mM: 189 sucrose, 26 NaHCO_3_, 10 D-glucose, 5 MgSO_4_, 3 KCl, 1.25 NaH_2_PO_4_, 0.2 CaCl_2_) bubbled with 95 % O_2_/5 % CO_2_. The brain was sectioned at 11° using a custom brain matrix as previously described (Banks et al 2015) and modified coronal slices were cut at 350 µm thickness using a vibratome (7000smz-2, Camden Instruments), hemisected and incubated at 34 °C for 1 hour after dissection in a slice holding chamber filled with artificial cerebrospinal fluid (aCSF, in mM: 124 NaCl, 26 NaHCO_3_, 10 D-glucose, 3 KCl, 2 CaCl_2_, 1.25 NaH_2_PO_4_, 1 MgSO_4_). Slices were subsequently stored at room temperature until use.

### Electrophysiology

Slices at ∼ 3.0 mm anterior to bregma were used for electrophysiology, placed in a submerged recording chamber and perfused with 34 °C aCSF at ∼2ml.min^-1^. A stimulating electrode (FH-Co, CBABAP50, USA) was placed on the hippocampal fibre bundle as previously described (Banks et al 2015). Pyramidal neurons in prelimbic cortex (layer 5 unless otherwise stated) were targeted under oblique infra-red illumination based on somatic morphology and patch clamped using 2-6 MΩ boroscillicate glass electrodes (GC150-10F, Harvard Apparatus) filled with potassium gluconate internal for current-clamp experiments (in mM: 120 k-gluconate, 40 HEPES, 10 KCl, 2 NaCl, 2 MgATP, 1 MgCl, 0.3 NaGTP, 0.2 EGTA, 0.1 Alexa-594 hydrazide) or cesium methylsulfonate for voltage clamp (130 CsMeSO_4_, 10 HEPES, 8 NaCl, 5 QX-314Cl, 4 MgATP, 0.5 EGTA, 0.3 NaGTP, 0.1 Alexa-594 hydrazide). Wide-field fluorescence was used at the end of experiments to confirm pyramidal cell morphology (prominent apical dendrite extending toward layer 1). Recordings were obtained using a Molecular Devices Multiclamp 700A or 700B, filtered at 4 KHz and digitized at a sample frequency ≥ 20 KHz with WinLTP2.30 (Anderson & Collingridge 2007) or pClamp10 software.

For current clamp recordings resting membrane potential (RMP) was recorded immediately after entering the whole-cell configuration. Intrinsic membrane properties were recorded as previously described (Barker et al 2017). Neurons were kept at −70 mV by injection of constant current throughout experiments. Synaptic stimulation occurred every 10 s, alternating between optogenetic stimulation of NRe and electrical stimulation of HPC input to achieve a 0.05 Hz intra-pathway basal stimulation frequency. Only neurons in which both pathways were measurable were used for synaptic experiments, this constituted 68% of neurons. Optogenetic stimulation was applied with a 470 nm LED (M470L3, Thorlabs) triggered by TTL pulses sent to an LEDD1B driver (Thorlabs) directed onto the soma and proximal dendrites via a 40x immersion objective (Olympus LUMPLFLN40XW) resulting in a 660 µm diameter illumination. The LED was usually driven at maximal strength, resulting in 4.35 mW.mm^-2^ light density. EPSP amplitude was adjusted by changing the light pulse duration (0.2-5ms, typically 1ms). The hippocampal fibre bundle was stimulated (0.1 ms duration) using a bipolar concentric stimulation electrode (CBAPB125, FHC) and Digitimer DS3 constant current stimulator. Stimulation intensity of each pathway was adjusted to achieve subthreshold EPSPs that were typically 2-8 mV.

NMDAR:AMPAR ratios were obtained by recording AMPAR-mediated synaptic potentials at –70 mV and, following bath application of 5 µM NBQX and 50 µM picrotoxin, NMDARs at +40 mV. Ro 25-6981 experiments were conducted at –40 mV to minimise cell death during experiments. NMDA receptor weighted decay time constants (τ_w_) were calculated in pClamp by fitting a double exponential curve between 90 - 10% of +40 mV NMDAR responses using the formula τ_w_ = τ1*A1/(A1+A2) + τ2*A2/(A1+A2) (Matta et al 2011). For MK-801 experiments, isolated NMDA receptor currents were obtained at +40 mV, 40 µM (+)MK-801 malleate was then bath applied for 10 minutes in the absence of stimulation with the cell held at −70 mV. Cells were then held at +40 mV again before resumption of stimulation at 0.1 Hz. For pairing plasticity experiments a baseline of 5 minutes was achieved within 10 minutes of break-in or cells were discarded to prevent washout (Malinow & Tsien 1990), intrinsic membrane properties were thus not recorded in those experiments.

### Staining and imaging

Rats were anaesthetised with Euthatal and perfused transcardially with phosphate buffer (PB) followed by 4% paraformaldehyde (PFA). Brains were removed and post-fixed in 4% PFA for 2 h before being transferred to 30% sucrose in PB for 48 h. Coronal sections (40 μm) were cut on a cryostat. Sections were washed three times in phosphate buffered saline (PBS), incubated for 30 min in 1% H_2_O_2_ in PBS, washed three times in 0.2 % Triton X in PBS (PBS-T), incubated for 1 h in blocking solution (5 % Normal goat serum, 2.5 % BSA, in PBS-T), then incubated in primary antibody solution (anti-mCherry rabbit polyclonal (Abcam ab167453); 1:1000 in blocking solution) for 24 h at RT. Sections were then washed four times in PBS-T, incubated in secondary antibody solution (goat anti-rabbit conjugated to Alexa Fluor 594 (Abcam ab150080); 1:500 in blocking solution) for 2 h at RT and then washed a further four times in PBS-T, mounted and coverslipped with Vectashield with DAPI (Vector Laboratories H-1500) mounting medium. Images were acquired using a widefield fluorescence microscope (Leica).

### Analysis and Statistics

Synaptic responses were averaged into one-minute bins and analysed using WinLTP2.3 or pClamp10. Intrinsic membrane properties were imported to MATLAB using SourceForge and analysed using code supplied by Dr Jon Brown (Exeter University, UK) as described previously (Barker et al 2017). Experiments used N numbers typical of the field, typically one cell per animal, number of cells was thus considered as experimental N, numbers of both cells and animals are reported. Data are expressed as mean ± SEM unless otherwise stated. Statistical tests were done using SPSS (IBM) or Graphpad Prism 7. Data sets were tested for normality prior to analysis using Shapiro-Wilk test to determine use of parametric or non-parametric tests as follows: unpaired observations were compared using student’s t-test or non-parametric Mann-Whitney U (two-tailed), unpaired observations of more than two groups were made using one-way ANOVA or non-parametric Kruskal Wallis with Tukey’s or Dunn’s post-hoc comparisons, respectively. Paired comparisons of two groups were made with paired t-test or Wilcoxon matched-pair signed rank (two-tailed). Comparisons across a range of within- and between-subjects variables were made with two-way repeated-measures ANOVA with Sidak’s post-hoc multiple comparisons. Short-term plasticity was assessed from normalised peak amplitudes with repeated-measures ANOVA with Greenhouse-Geisser corrections applied where Maulchy’s test for Sphericity significance was < 0.05. For pairing experiments, plasticity in individual pathways was assessed by comparison of average EPSP amplitudes during baseline and 30-40 minutes post-pairing (follow-up). For all experiments, significance was reported at p < 0.05.

## Results

### Characterising L5 pyramidal neurons in prelimbic cortex that receive nucleus reuniens and hippocampal inputs

Of the thalamic nuclei with projections to mPFC, viral injections into ventral midline thalamus transduced reuniens, ventral reuniens and rhomboid nuclei (collectively referred to as NRe) with channelrhodopsin variant ChETA_TC_ (AAV9-CaMKii-hChR2(E123T/T159C)-mCherry; Fig 1A) resulting in mCherry labelling (Fig 1B) across all layers (1-6) of mPFC (Vertes et al 2006). In this study we focussed on the NRe and HPC projections to layer 5 pyramidal neurons - one of the primary output cells of prelimbic cortex and the main target for HPC afferents (Liu & Carter 2018). Modified coronal mPFC slices were prepared as described previously (Banks et al 2015, Parent et al 2010). To activate NRe axons in PFC an LED was directed over L5 cell somata and proximal dendrites via the microscope objective (Fig 1C). To activate the HPC fibre bundle, a stimulating electrode was positioned between the dorsal tenia tecta and the nucleus accumbens (Fig 1C; Banks et al 2015). This approach allowed us to simultaneously compare light evoked NRe and electrically evoked HPC inputs onto the same cells in PFC. ChETA_TC_ activation of NRe afferents resulted in EPSPs with simple waveforms providing direct evidence that neurons of NRe synapse upon L5 pyramidal cells in mPFC (Fig 1D). Analysis of 187 cells showed there is a high degree of convergence of HPC and NRe pathways onto individual mPFC pyramidal cells; 68 % of L5 pyramidal neurons received input from both NRe and HPC, 18 % from HPC alone and 7 % from NRe alone (Fig 1E).

Layer 5 pyramidal neurons can be separated by their projection targets: intratelencephalic (IT) which project cortically, and pyramidal tract (PT) which project subcortically; these classes of neuron have distinct electrophysiological properties (Dembrow et al 2010). To determine whether HPC or NRe afferents differentially target IT or PT neurons we compared intrinsic passive and active membrane properties of cells receiving input from both NRe and HPC, either pathway alone, or neither input. Cells which only had input from HPC had significantly lower median input resistance compared to cells which received both NRe and HPC input (Fig 1E). Cells with only HPC input also expressed the largest percentage of I_h_-mediated voltage sag; however there was no difference in the sum of absolute level of sag and the rebound from hyperpolarisation, a measure which has been used to classify cortically and subcortically projecting cells previously (Gee et al 2012, Lee et al 2014). Resting membrane potential and membrane time constant were the same across all input groups (Fig 1E).

Regarding active membrane properties, action-potential threshold of cells receiving only HPC input was more hyperpolarised than that of either group of neurons which received NRe input (Fig S1A). No other difference in action potential properties (Fig S1A), number of spikes or spike frequency adaptation was found between groups (Fig S1B). Amplitudes of afterhyperpolarisation and afterdepolarisation were not different across groups (Fig S1C).

These results show that cells receiving only NRe- or NRe and HPC input have higher input resistance and less sag than those receiving only HPC input. These data suggest that layer V pyramidal cells receiving NRe and NRe/HPC inputs have intrinsic membrane properties consistent with an IT phenotype. HPC input has been suggested to preferentially target IT cells, however in this study those cells in which only HPC input is present appear to be of PT type (Anastasiades et al 2018, Dembrow et al 2010, Dembrow et al 2015). Subsequent experiments were performed only in cells with both NRe and HPC inputs.

### Comparison of nucleus reuniens and hippocampal synaptic properties

Activation of NRe and HPC afferents resulted in simple waveform EPSPs which were highly alike (Fig 2A). Bath application of tetrodotoxin (TTX, 0.5 µM) abolished NRe (and HPC) EPSPs (Fig S1D,E) confirming that ChETA_TC_-evoked NRe responses are action potential dependent. Addition of voltage-dependent K^+^ blocker 4-AP (100 µM) partially restored NRe transmission when stimulus duration was increased (Fig S1D,E), demonstrating NRe EPSPs onto L5 pyramidal neurons are monosynaptic (Petreanu et al 2009). In further support of the monosynaptic nature of NRe EPSPs under control conditions, ChETA_TC_-EPSPs were of short latency, being moderately faster than, but not significantly different to those evoked electrically from HPC fibres (Fig 2B, Mann-Whitney p = 0.077, n =33 cells from 28 animals).

**Figure 2.**
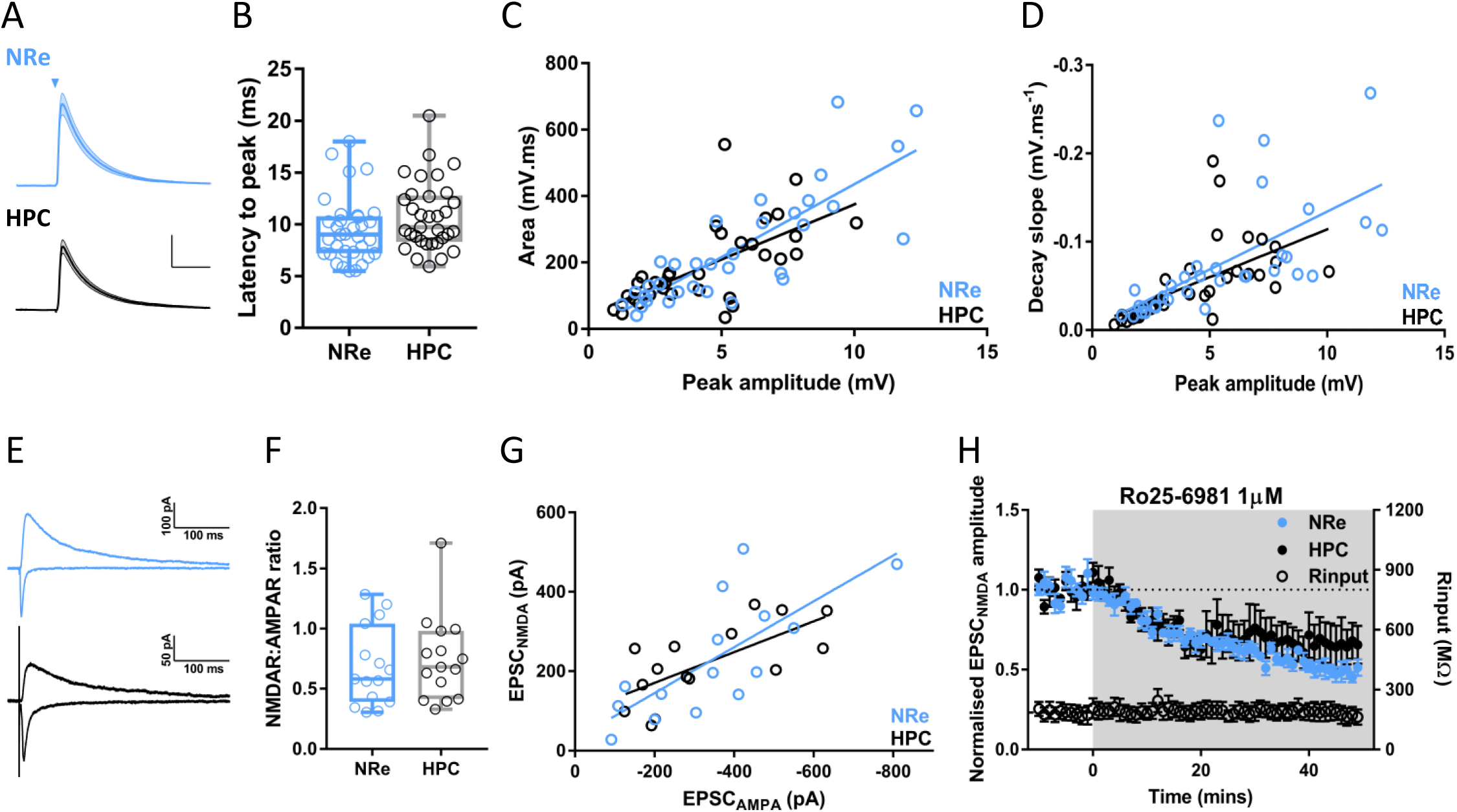
NRe and HPC synapses are indistinguishable. **A –** Average waveforms of NRe and HPC EPSPs, stimulation denoted by triangle, electrical stimulation artefacts removed for clarity. Traces show mean ± SEM waveform, scale bars = 2 mV/50 ms. **B -** Latency from stimulation to EPSP peak, box plot shows median, 25 and 75^th^ percentiles, whiskers maxima and minima. Individual values shown as open circles. (Mann-Whitney test, U = 407, p = 0.077, n = 33 cells from 28 animals,). **C** – EPSP peak amplitude vs area of NRe and HPC EPSPs. Linear regression slopes were not significantly different (F_(1,62)_ = 1.5, p = 0.23). **D** – EPSP peak amplitude vs decay slope from 90-15 % of EPSP peak. Linear regression slopes were not significantly different (F_(1,62)_ = 0.3, p = 0.57). **E** – Representative −70 mV AMPAR-mediated (negative-going) and +40 mV NMDAR-mediated (positive-going) EPSCs resulting from NRe (blue) and HPC (black) stimulation. **F** – NMDAR:AMPAR ratios of NRe and HPC EPSCs were not significantly different (Mann-Whitney U = 99, p = 0.60, n = 15 cells from 10 animals). **G** – EPSC_AMPA_ vs EPSC_NMDA_ for NRe and HPC inputs. Linear regression slopes were not significantly different (F_(1,26)_ =1.05, p = 0.32). **H** – EPSC_NMDA_ inhibition by bath application of GluN2B selective antagonist Ro25-6981 (3 µM) as indicated by grey shaded region. Mann-Whitney U = 12, p = 0.4. N = 6 cells from 5 animals.

Bath application of ionotropic glutamate receptor antagonists NBQX and D-AP5 completely blocked both NRe (FigS1D,E) and HPC responses (Fig S1F), demonstrating that NRe input is glutamatergic (Hur & Zaborszky 2005). Linear regression of EPSP amplitude vs area for NRe and HPC yielded equal slopes (Fig 2C, p = 0.23) and regression of EPSP amplitude vs decay slope was also indistinguishable (Fig 2D, p = 0.57). Since NRe and HPC EPSPs are recorded within the same cells, there can be no difference in intrinsic electrical properties between cells receiving these synaptic pathways. This suggests that there is little difference in AMPA receptor subunit composition between synapses, and that the dendritic location of activated synapses is similar (Magee 2000).

NMDA receptors, in particular GluN2B subunit containing receptors, play a key role in supporting persistent firing, which is thought to contribute to working memory in PFC (Monaco et al 2015), moreover expression of GluN2B subunits may be input specific (Flores-Barrera et al 2014). We therefore examined NMDA receptor expression and stoichiometry at NRe and HPC synapses. No difference in the ratio of NMDAR:AMPAR EPSCs was seen (Fig 2E,F; Mann-Whitney, p = 0.60, n = 15 cells from 10 animals) and linear regression of EPSP_AMPA_ vs EPSC_NMDA_ slopes were equal (Fig 2G, p = 0.32). Weighted decay time constants were not significantly different (Tw: NRe = 135.3 ± 44.3 ms, HPC = 128.9 ± 32.8 ms, Mann Whitney U = 34, p = 0.9) and inhibition of GluN2B subunit-containing receptors by Ro 25-6981 (3 µM) was indistinguishable between the two inputs (Fig 2H, Mann-Whitney, p = 0.4, n = 6 cells from 5 animals), indicating that overall levels of NMDAR expression and stoichiometry are not appreciably different at NRe and HPC synapses. Altogether these data show that single evoked NRe and HPC EPSCs in prelimbic cortex L5 pyramidal cells are indistinguishable and likely to have similar postsynaptic ionotropic glutamate receptor composition.

### Nucleus reuniens, but not HPC, afferents to mPFC undergo short-term depression at theta frequency

Next we examined short-term plasticity of NRe and HPC inputs to L5 pyramidal neurons at frequencies relevant to NRe-mPFC and HPC-mPFC theta (4-12 Hz) oscillations (Hallock et al 2016, Jones & Wilson 2005, O’Neill et al 2013, Roy et al 2017, Siapas et al 2005, Spellman et al 2015) and the tonic firing frequency of NRe matrix cells (Walsh et al 2017). In contrast to *in vivo* recordings showing paired-pulse facilitation (Di Prisco & Vertes 2006), NRe EPSPs strongly depressed when activated at both 5 and 10 Hz (Fig 3A), an effect we did not see for HPC EPSPs (Liu & Carter 2018). Pronounced short-term depression of NRe responses was also observed in L2/3 pyramidal cells, with no difference in depression seen between layers 2/3 and 5 (Fig 3B). These data demonstrate a specific functional projection from NRe to L2/3 pyramidal neurons and show that, in contrast to the HPC projection, NRe projections to mPFC pyramidal cells undergo short term depression irrespective of mPFC lamina.

**Figure 3.**
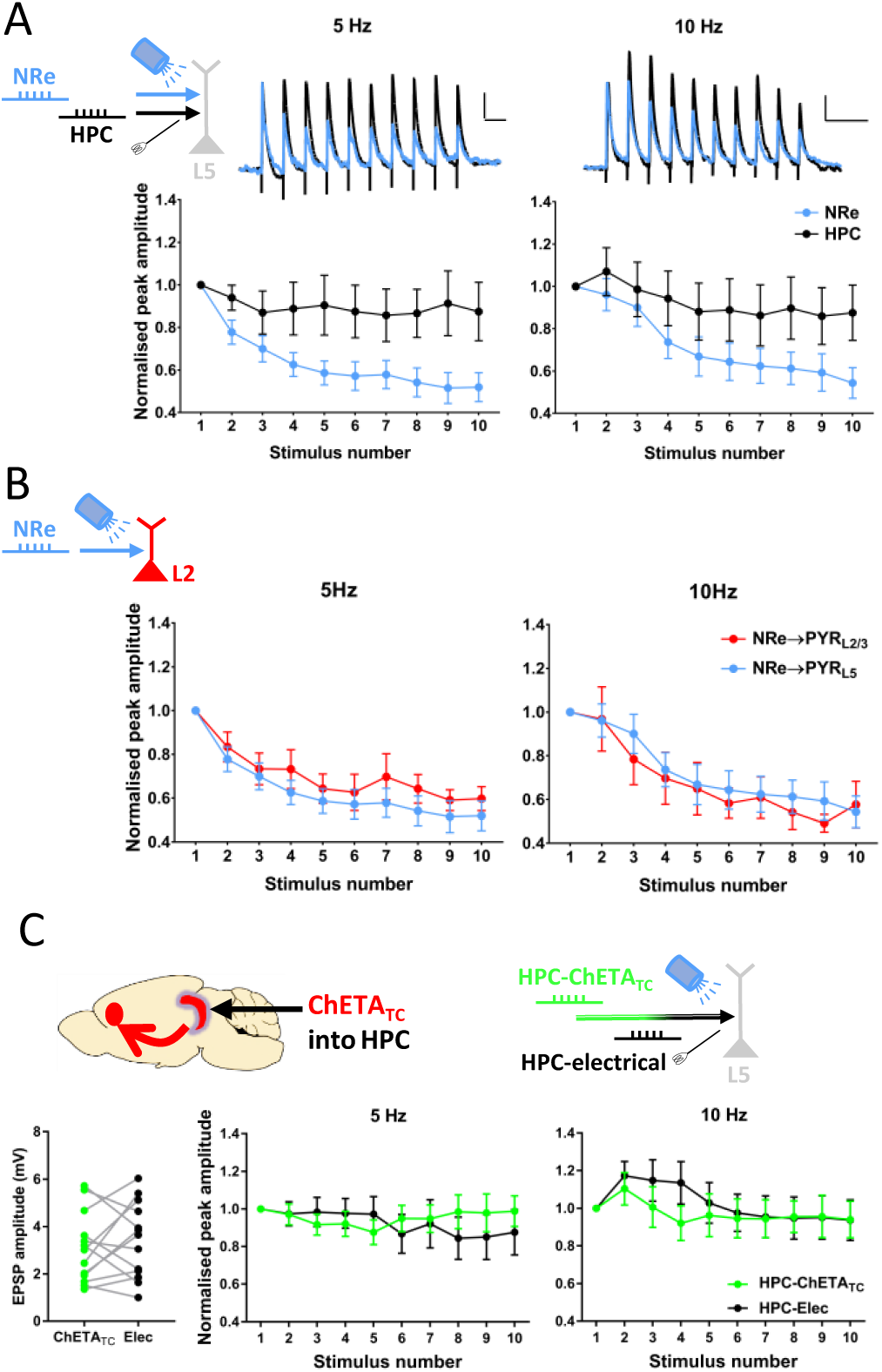
Nucleus reuniens input to prelimbic cortex depresses at theta frequency. **A** - NRe inputs to L5 pyramidal neurons undergo strong short-term depression, and show different plasticity pattern to HPC inputs at 5Hz (repeated measures two-way ANOVA: main effect of pathway F_(1,11)_ = 8.5, p = 0.014; main effect of response number F_(1.9,20.6)_ = 5.1; p = 0.018, interaction F_(3.1, 34.1)_=4.4, p = 0.0095)and 10 Hz (pathway F_(1,11)_ = 5.0, p = 0.048; response number F_(3.2,35.4)_ = 27.9, p = 1.1 × 10^−9^; interaction F_(4.3,47.0)_ = 8.5, p=0.00002; Greenhouse-Geisser correction applied, both frequencies). N= 12 cells from 11 animals. **B** – NRe inputs to L2/3 pyramidal cells show equal degree of short-term depression as inputs to L5 at 5 Hz (Repeated-measures two-way ANOVA: main effect of layer F_(1,19)_ = 0.71, p = 0.41; main effect of response number F_(8,61.0)_ = 15.3, p = 2.6 × 10^−16^; interaction F_(3.2,61)_ = 0.45, p = 0.73) and 10Hz (main effect of layer: F_(1,19)_ = 0.12, p = 0.73; main effect of response number F_(8,62.7)_ = 30.3, p = 6.8 × 10^−28^; interaction F_(3.3,62.7)_ = 0.9, p = 0.44; Greenhouse-Geisser correction applied, both frequencies. L2/3 n = 9 cells from 5 animals; L5 data repeated from Fig 3A). **C** – Following injection of AAV9-CaMKii-hChETATC-mCherry into intermediate/ventral HPC, transmission evoked by electrical and optogenetic stimulation were compared. EPSPs evoked by ChETA_TC_ and electrical stimulation were of similar amplitude (paired t-test, t_(12)_ = 0.78, p = 0.45). No difference in short-term plasticity was observed at 5 (main effect of stimulation method 5 Hz: F_(1,12)_ = 0.07, p = 0.79; main effect of response number F_(2.1,30.0)_ = 0.45, p = 0.65; interaction F_(3.1,36.9)_ = 2.2, p = 0.10) or 10 Hz stimulation frequency (stimulation method F_(1,12)_ = 0.38, p = 0.55; response number F_(2.1,24.7)_ = 4.6, p = 0.02, interaction F_(2.6,31.7)_ = 1.1, p =0.38). Greenhouse-Geisser corrections applied. N = 13 cells from 5 animals.

We first addressed the possibility that short-term depression of NRe inputs was an artefact of optogenetic activation since ChR2 variants might be unable to evoke spiking reliably at higher frequencies (Berndt et al 2011) and transduction of neurons with AAVs itself may affect synaptic release (Jackman et al 2014). To test this possibility, we transduced the HPC with ChETA_TC_ and compared, in the same mPFC cells, HPC EPSPs evoked by electrical and optical stimulation. Electrical and ChETA_TC_ evoked EPSPs were of similar initial amplitude and no significant difference was observed between optical and electrical stimulation of HPC afferents at theta frequencies (Fig 3C), thereby demonstrating the ability of ChETA_TC_ to evoke high fidelity transmission at 5 and 10 Hz. We also compared the degree of short-term depression observed in NRe inputs to the duration of light pulses used to evoke NRe EPSPs, with no relationship observed between these two variables (supplementary S2A). These data demonstrate that short-term depression of NRe transmission is not an artefact of optogenetic activation of NRe inputs.

For completeness we also tested in hippocampal afferents the range of stimulation frequency possible with ChETA_TC_. At frequencies of 20 Hz and above we saw deficits in optically-evoked HPC transmission compared to electrically evoked transmission (Fig S2B), with optical EPSPs showing attenuated amplitude at 20 Hz and an inability to evoke subsequent EPSPs at 100 Hz, possibly owing to failure to recover from desensitisation between pulses.

Having demonstrated that pronounced short-term depression is a physiological feature of NRe-mPFC synapses we set about exploring possible mechanisms for this depression. One potential mechanism for synaptic depression is activation of presynaptic metabotropic glutamate receptors (mGluRs) leaving to autoinhibition. Since activation of group II mGluRs depresses MD inputs to mPFC (Joffe et al 2020) and mGluR2 has been shown to be strongly expressed in reuniens and rhomboid nuclei (Lourenco Neto et al 2000, Ohishi et al 1998), we therefore hypothesised that differential expression of group II mGluRs may underlie the differences in short-term depression occurring at NRe and HPC inputs. However, application of the Group II mGluR agonist LY354740 (500 nM) resulted in similar levels of acute depression of both NRe and HPC inputs (Fig 4A). Furthermore, neither the broad spectrum mGluR antagonist LY341495 (100 µM) nor the selective group II mGluR antagonist EGLU (10 µM) affected the short-term dynamics at 5 or 10 Hz in either pathway (Fig 4B; S3A).

**Figure 4.**
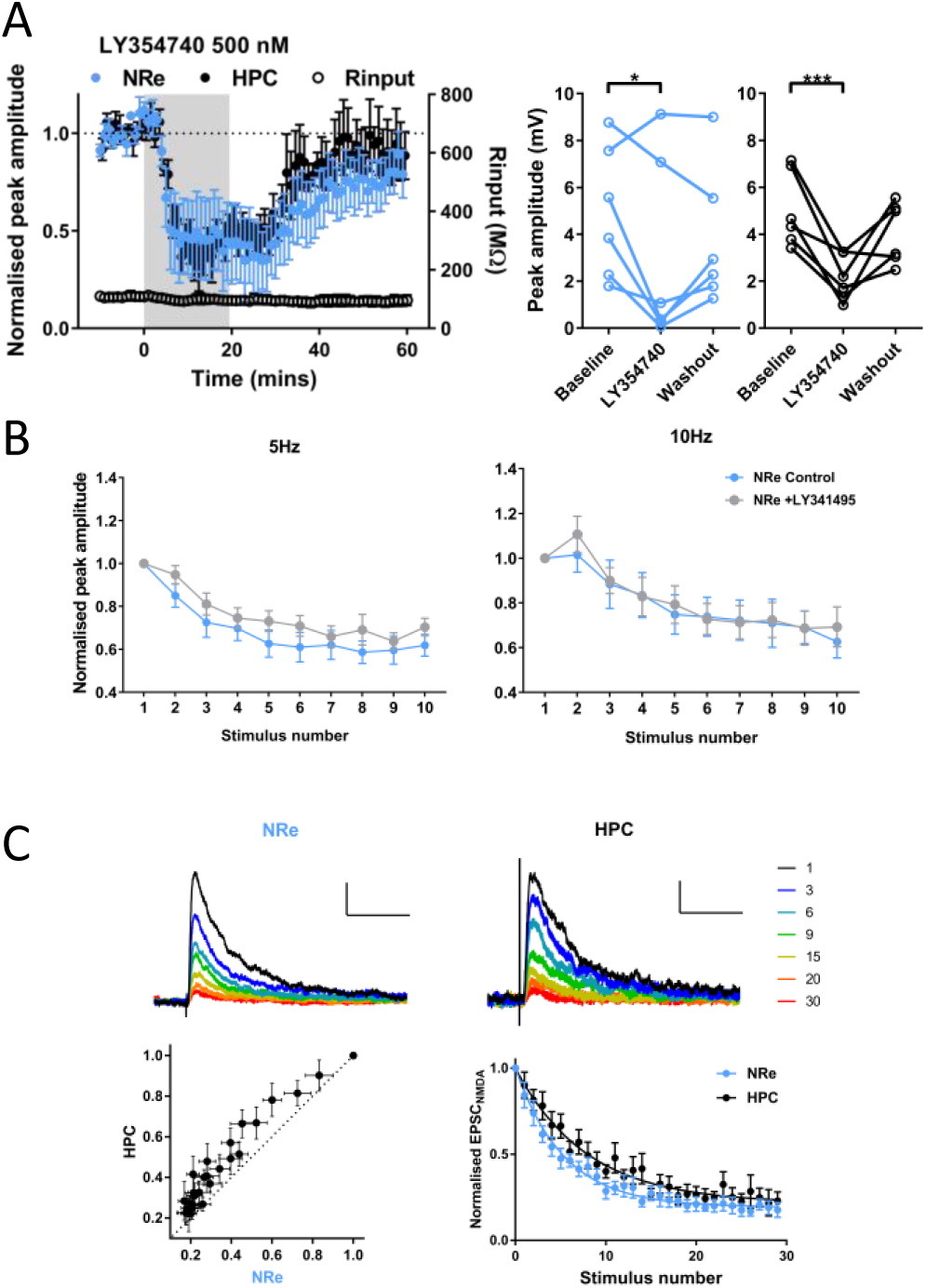
NRe and HPC are inhibited by group II mGluR activation, but high probability of release underlies short-term depression of NRe inputs. **A -** Activation of group II mGluRs with LY354740 (500 nM) reveals no difference in acute depression of NRe and HPC inputs measured during the final 10 minutes of drug application (two-way repeated measures ANOVA:main effect of timepoint F_(2,20)_ = 13.3, p = 0.0002; pathway F_(1,10)_ = 0.02, p = 0.90; interaction F_(2,20)_ = 0.81, p = 0.46; Sidak’s post-hoc comparisons shows difference vs baseline, NRe p= 0.024, HPC p = 0.0009). N = 7 cells from 7 animals. **B** – LY341495 (100 µM) did not affect short-term plasticity of NRe input at 5 or 10 Hz (RM ANOVA; 5Hz: main effect of drug F_(1,8)_ = 4.8, p = 0.059; main effect of response number F(4.0,32.3) = 23.6, p = 3 × 10^−9^; interaction^#^ F_(4.4,35.2)_ = 0.85, p = 0.51; 10Hz: main effect of drug F_(1,8)_ = 0.47, p = 0.51; response number F(2.6,20.7) = 28.8, p = 2.9 × 10^−7^; interaction F_(2.5,20.2)_ = 2.6, p = 0.089). Greenhouse-Geisser corrections applied. N= 9 cells from 6 animals. **C** – Activity-dependent block of isolated NMDA EPSCs by MK-801 (40 µM). Example blockade of EPSC_NMDA_ measured at +40 mV in NRe and HPC pathway, traces coloured by stimulus number in presence of MK-801, normalised to amplitude of first response, scale bars: 0.25 of normalised peak/100 ms. Plot of NRe vs HPC amplitudes shows data lie above the identity line. Decay of NRe is significantly faster than decay of HPC (single exponential curve constrained to Y0 = 1, NRe τ = 5.4, HPC τ = 7.9, extra sum of squares F-test, F_(2,532)_ = 19.0, p < 0.0001). N= 10 cells from 8 animals.

We next explored the possibility that recruitment of a G-protein coupled potassium conductance responsible for NRe depression. Theta range stimuli were thus delivered before and after application of the GABA_B_ receptor antagonist CGP55845 (1 µM), however no effect was seen on either pathway (Fig S3B). Additionally, neither the NMDAR antagonist D-AP5 (50 µM; Fig S3C) nor the nicotinic receptor antagonist mecamylamine (1 µM; Fig S3D) affected short term plasticity, suggesting no role for presynaptic NMDARs or nAChRs in regulation of synaptic release.

Given the above results we hypothesised that presynaptic release mechanisms in NRe synapses most likely explain short-term depression, possibly owing to a high initial release probability as has been shown for other thalamocortical synapses (Gil et al 1999). To address this question, we measured the rate of blockade of pharmacologically isolated NMDAR-mediated currents (EPSC_NMDA_) by the use-dependent NMDA receptor antagonist MK-801 (Gil et al 1999, Hessler et al 1993). As MK-801 blocks only open channel pores, pathways with high release probability are predicted to activate NMDA receptors at a large proportion of their synaptic sites, thus resulting in a faster rate of block by MK-801 compared to a pathway with lower release probability. NRe EPSC_NMDA_ were blocked faster than HPC EPSC_NMDA_ as shown by plotting EPSC_NMDA_ amplitudes of NRe and HPC for each trial against each other, with the data points lying above the line of identity (Fig 4C). Furthermore, decay curves of the time course of block by MK-801 were fit by single exponential curves with significantly different parameters (p < 0.0001). Together these data show that NRe has a high probability of release, as observed at other thalamocortical synapses, and this is likely to underlie the marked differences between NRe and HPC transmission observed during theta-frequency stimulation.

### Cholinergic neuromodulation of HPC, but not NRe afferents via M2 muscarinic receptors

Acetylcholine signalling in mPFC is essential for associative recognition memory (Barker & Warburton 2009) and spatial working memory (Ragozzino & Kesner 1998), and additionally phasic ACh release in mPFC is associated with high cognitive load, cue-detection and promotion of HPC-mPFC coherence (Howe et al 2017, Teles-Grilo Ruivo et al 2017). Given the importance of NRe function in these behaviours and in regulation of oscillatory activity (Barker & Warburton 2018, Hallock et al 2016) it is likely that NRe inputs to mPFC are active during phasic ACh release in mPFC, however it is unknown whether NRe afferents are sensitive to modulation by ACh. To address this question, we bath applied the broad-spectrum cholinergic agonist carbachol (CCh) for 10 minutes. To our surprise, NRe afferent input to L5 pyramidal neurons was unaffected by CCh (Figs 5A, S4A). In contrast, micromolar concentrations of CCh produced a strong, reversible attenuation of HPC inputs (Fig 5A, individual experiments shown in Fig S4A; 2-way repeated measures ANOVA pathway x concentration x timepoint interaction of CCh effects (F_(4,40)_ = 3.0, p = 0.029).

**Figure 5.**
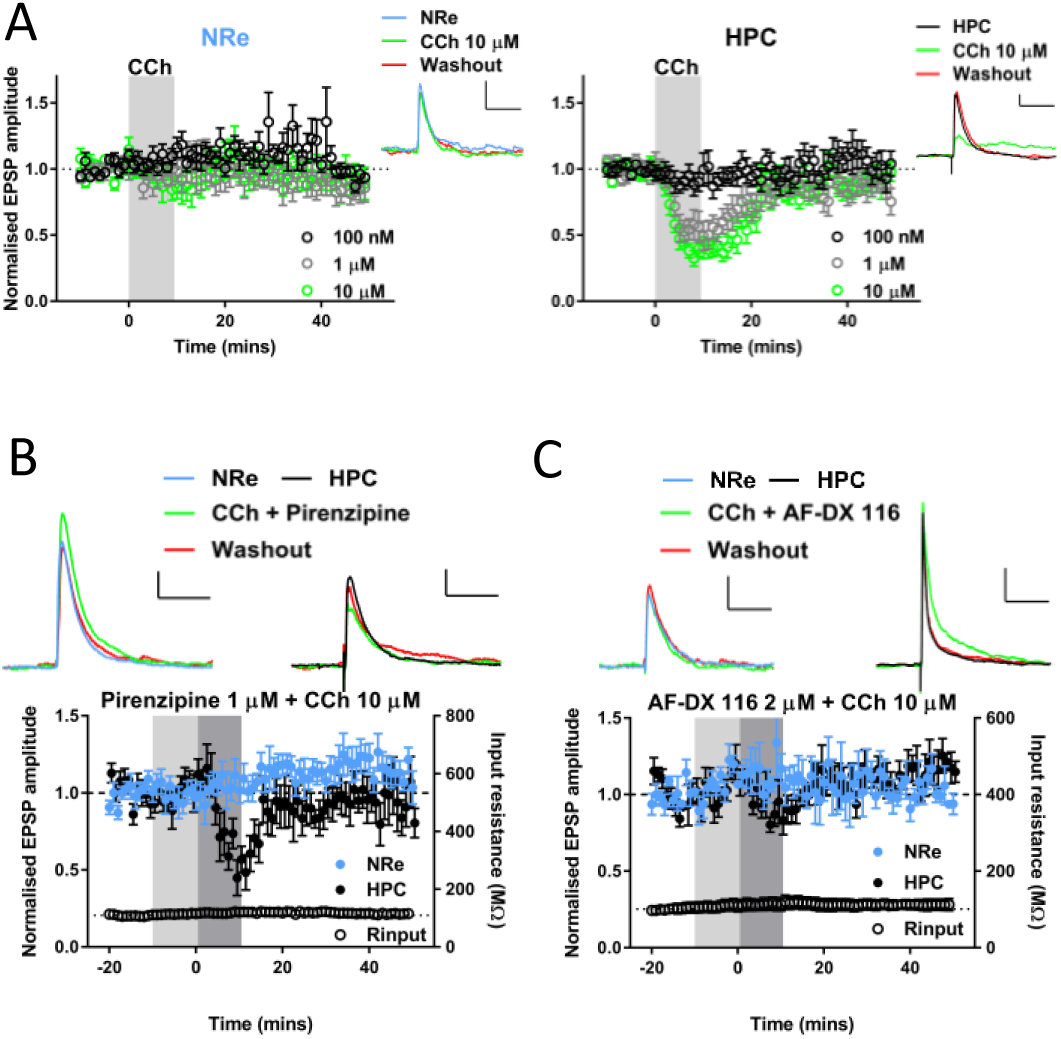
Cholinergic modulation of hippocampal, but not nucleus reuniens inputs to prelimbic cortex via M2 muscarinic receptors. **A** – Pooled data showing 10-minute bath application of cholinergic agonist carbachol (shaded region) at different concentrations has no effect upon NRe (left) input to PrL, but reversibly depresses HPC inputs (right) in a concentration-dependent manner. Example EPSPs for each pathway at baseline (−10 to -1 mins), acute (10-19 mins) and washout (40-49 mins) shown from a representative 10 µM experiment (scale bars = 1 mV, 100ms; n = cells/animals: 100 nM = 9/8; 1 µM = 6/4; 10 µM = 8/7). **B**- Selective M1 muscarinic antagonist pirenzipine does not block depression of HPC by 10 µM CCh. Pirenzipine (1 µM) pre-applied during light shaded region and co-applied with CCh during dark shaded region. EPSPs from a representative cell are shown above pooled data, scale bars = 1 mV/100 ms. N= 7 cells from 6 animals. **C** – Selective M2 muscarinic antagonist AF-DX 116 blocks depression of HPC by 10 µM CCh. AF-DX 116 (2 µM) pre-applied during light shaded region and co-applied with CCh during dark shaded region. EPSPs from a representative cell are shown above pooled data, scale bars = 1 mV/100 ms for NRe and 2 mV/100 ms HPC. N = 5 cells from 5 animals. For plots of individual experiments and statistics please refer to Fig S4.

We next sought to characterise the receptors responsible for the depression of HPC afferents by CCh, focussing on muscarinic receptors as these have previously been shown to mediate CCh depression of various synapses in mPFC (Caruana et al 2011, Huang & Hsu 2010, Wang & Yuan 2009). The M1 selective muscarinic antagonist pirenzepine (1 µM) did not prevent the HPC-mPFC CCh depression (Figs 5B, S4B). In contrast, the M2 receptor antagonist AF-DX 116 (2 µM) completely blocked the acute depression of HPC responses by CCh (p = 0.001; Fig5C, S4B). NRe inputs were not significantly modulated by CCh in the presence of M1 or M2 selective antagonists, thus ruling out the possibility that M1 and M2 receptor activation have equal and opposite effects that mask each other (Figs 5B-C, S4B). Together these data suggest that during increased cholinergic tone, HPC inputs to L5 pyramidal cells in mPFC are selectively inhibited via M2 muscarinic receptors. In contrast the NRe-mPFC input, somewhat uniquely among studied mPFC glutamatergic synapses, is not strongly modulated by cholinergic activation.

### Dopaminergic neuromodulation of NRe and HPC afferents

Having shown that NRe and HPC inputs are differentially controlled by cholinergic neuromodulation, we wanted to determine whether input specific modulation was a common feature of these pathways. Dopaminergic signalling in the mPFC plays a key role in executive function in mPFC (Ott & Nieder 2019) and, amongst other effects, dopamine is known to modulate synaptic transmission in mPFC, with varied effects upon local and distal excitatory synapses (Gao et al 2001, Gonzalez-Islas & Hablitz 2003, Law-Tho et al 1994, Seamans et al 2001, Urban et al 2002). We therefore examined the effect of D1R and D2R dopamine receptor subclass agonists on basal NRe and HPC EPSPs.

The D1-like receptor agonist SKF81297 (0.5 µM) produced small increases in the amplitude of both inputs (Figs 6A, S5A) but a statistically significant difference was only achieved in the HPC pathway. 10 µM SKF81297 also enhanced responses in both inputs but this was statistically insignificant (Figs 6B, S5A) and there was no effect on paired-pulse ratio (100 ms inter-stimulus-interval) in either pathway (Fig S5B). The activation of D2R-like receptors by quinpirole (10 µM) did not modulate NRe or HPC inputs (Fig 6C, S5C).

**Figure 6.**
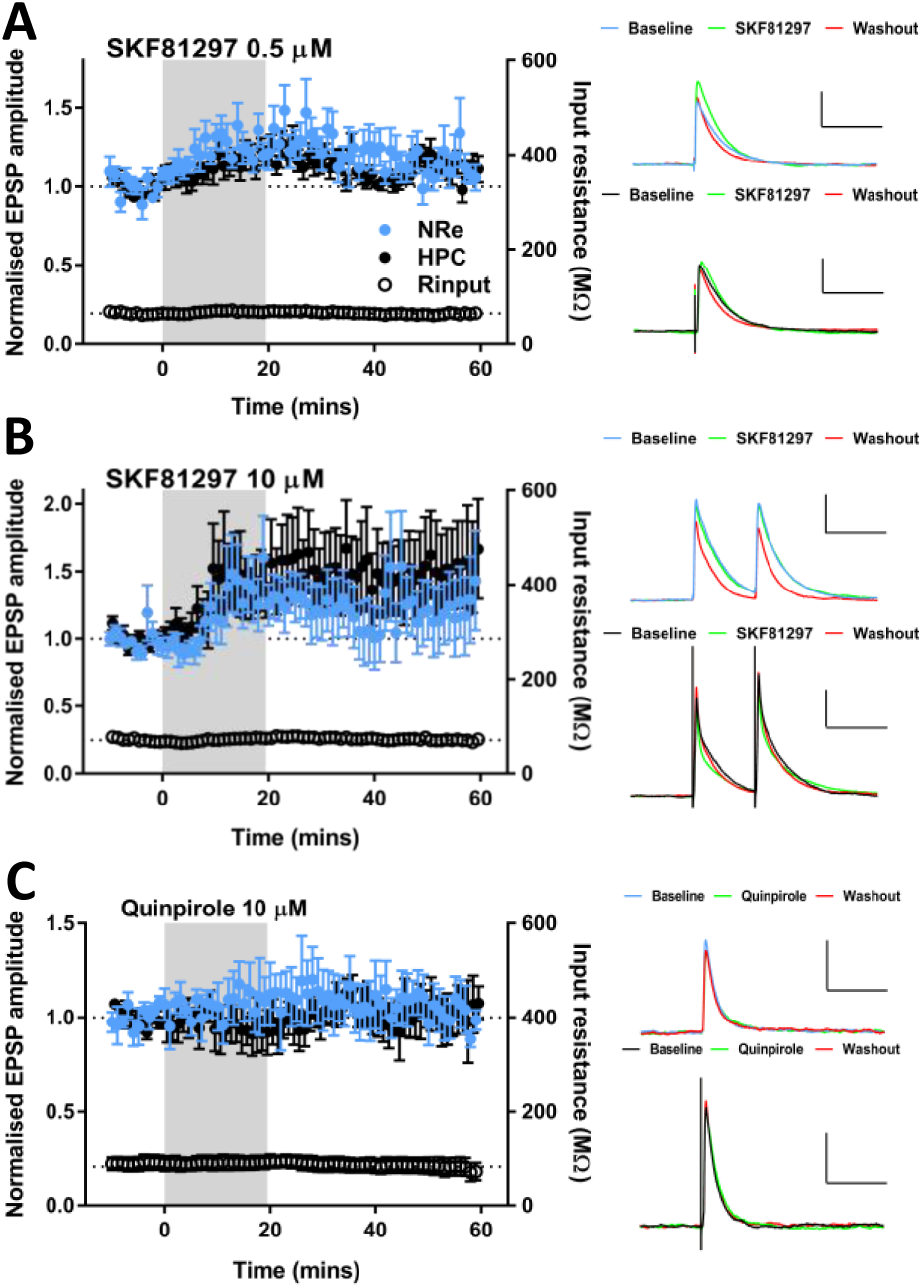
Dopaminergic modulation of basal NRe or HPC inputs to mPFC. **A** – D1R-like dopamine agonist SKF81297 bath applied at 0.5 µM caused a modest, reversible increase in transmission (two-way ANOVA, main effect of timepoint: F_(2,28)_ = 5.9, p = 0.0072; pathway F_(1,14)_ = 1.1, p = 0.31; interaction F_(2,28)_ =1.2, p = 0.31), Sidak’s post-hoc HPC baseline vs SKF81297 p = 0.013). Values = mean ± SEM. Representative EPSPs at baseline, final 10 mins of drug application and final ten mins of recording, scale bars = 3mV/100ms. N= 9 cells from 9 animals. **B** – SKF81297 at 10 µM did not result in a significant alteration of NRe or HPC EPSPs (two-way ANOVA, main effect of timepoint F_(2,26)_ = 1.9, p = 0.17; main effect of pathway F_(1,18)_ = 0.3, p = 0.59, interaction F_(2,36)_ = 0.55, p = 0.58). Values = mean ± SEM. Representative EPSPs shown, scale bars = 3mV/100ms. N= 10 cells from 8 animals. **C** –D2R-like dopamine agonist quinpirole (10 µM) does not affect basal NRe or HPC transmission (two-way ANOVA, main effect of timepoint F_(2,20)_ = 0.05, p = 0.31, interaction F_(2,20)_ = 1.2, p = 0.31). Data shown are mean ± SEM, representative EPSPs (scale bars = 2mV/100ms), raw EPSP amplitudes for individual experiments. N= 6 cells from 6 animals.

Previous reports have argued that dopamine receptor expression is tightly linked to intrinsic cellular properties, with L5 pyramidal neurons that express D2-like receptors showing prominent hyperpolarisation-activated current (I_h_) and D1-expressing neurons lacking prominent I_h_ (Gee et al 2012). However, we found no relationship between the magnitude of I_h_ and modulation by either D1R or D2R agonists (Fig S5D,E).

### NRe and HPC inputs undergo associative synaptic plasticity via NMDA receptor activation

Since both NRe and HPC inputs to mPFC are required for memory encoding we next asked whether NRe inputs to mPFC undergo synaptic plasticity and whether NRe and HPC inputs interact to induce synaptic plasticity. We hypothesised that a signal originating in either HPC or NRe would project directly to mPFC and di-synaptically to mPFC via the other region, thus resulting in a short lag between inputs (Fig 7A). We therefore paired stimulation of NRe and HPC afferents at time windows predicted from a simplified HPC-NRe-mPFC circuit (Dolleman-van der Weel et al 2019). HPC and NRe stimuli were paired with 10 ms inter-stimulus-intervals (ISIs). Comparable patterns of activity have been shown to induce so called input-timing dependent plasticity (ITDP) in the hippocampus (Dudman et al 2007). The pairing of HPC and NRe inputs was performed at 5 Hz since HPC-mPFC coherence in the theta range has been associated with performance of working memory tasks (Jones & Wilson 2005, Siapas et al 2005) and theta coherence can be enhanced following learning (Benchenane et al 2010). NRe plays a role in coordination of HPC-mPFC oscillations at theta (Hallock et al 2016, Kafetzopoulos et al 2018, though see: Roy et al 2017) and delta (2-5Hz; Roy et al 2017) and contains cells with spontaneous firing across these ranges (Walsh et al 2017).

**Figure 7.**
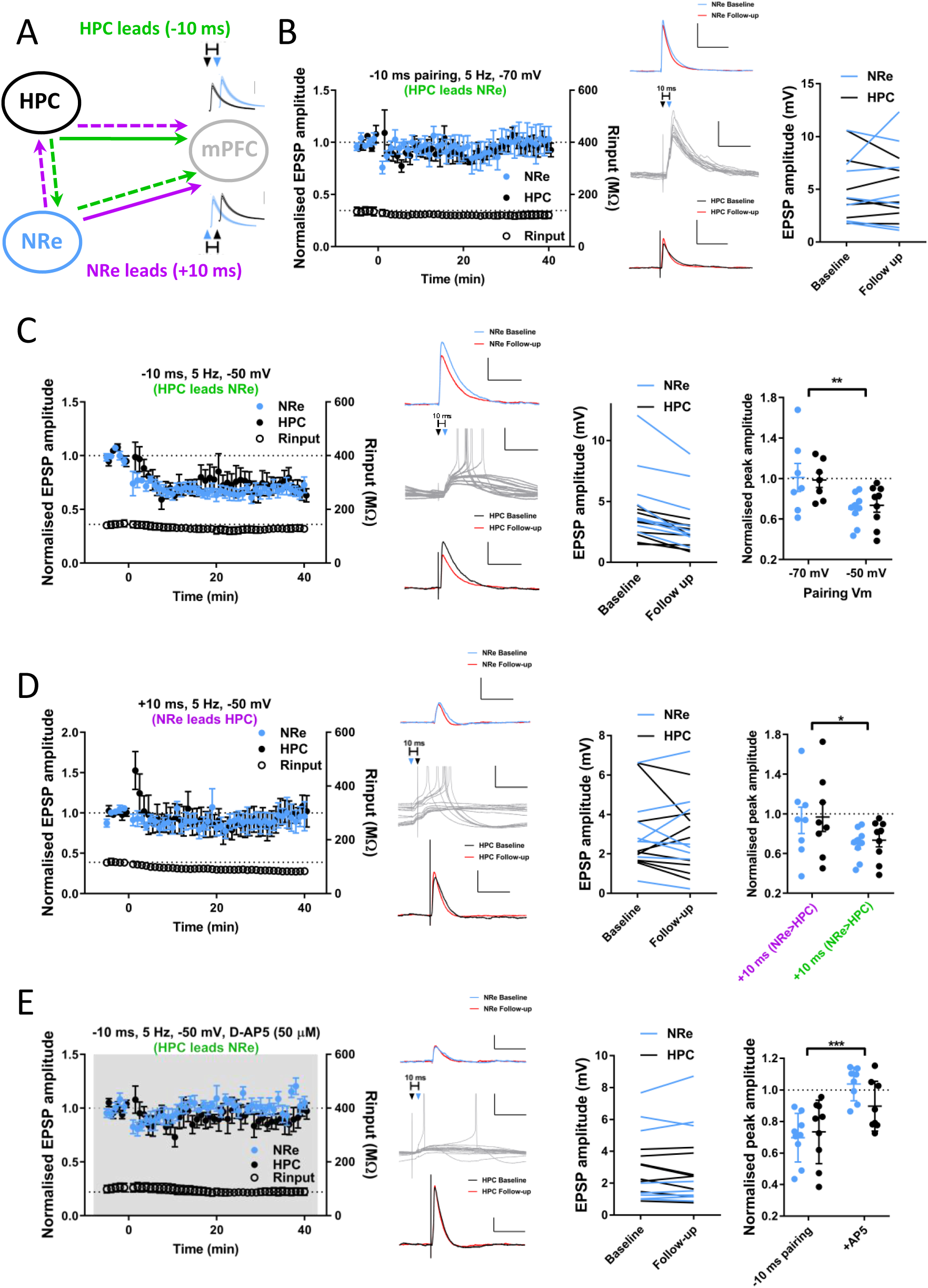
Pairing of HPC and NRe inputs repeated at theta frequency induces NMDA receptor-dependent, associative LTD. **A** – Schematic diagram showing hypothesised tripartite circuit dynamics. Information arising in HPC (green pathway) may project directly to mPFC (solid arrow) and feed forward disynaptically via NRe (dashed arrows), HPC EPSPs would therefore precede NRe in mPFC resulting in negative lag. Conversely a signal originating in NRe may reach mPFC directly and disynaptically via HPC (purple) resulting in the opposite temporal activation profile. **B** – Theta-frequency pairing of HPC and NRe inputs at −70 mV (HPC stimulus preceding NRe by 10ms, 100 pairs delivered at 5 Hz, Vm = −70 mV) does not induce synaptic plasticity in either pathway (Follow up: plasticity measured at 30-40 mins after pairing; paired t-test: NRe t_(6)_ = 0.6, p = 0.56; HPC t_(6)_ = 0.8, p = 0.45). Traces show example averaged EPSPs at baseline (NRe; blue (above)/HPC; black (below)) and 30-40 mins (follow-up; red) and the first 15 pairings (grey; middle traces) with HPC stimulation denoted by black and NRe by blue triangles, respectively. Scale bars EPSPs: 5 mV, 100 ms, pairing: 5 mV, 50 ms. Right: EPSP amplitudes for individual experiments. N = 7 cells from 7 animals. **C** – Depolarisation to −50 mV during 5 Hz pairing (−10 ms delay) induces LTD of NRe and HPC inputs to PFC. Traces as in B except scale = 2 mV in non-pairing traces. Example experiment traces show baseline and 30-40 min EPSPs. Note incidence of spiking during pairing protocol. Individual experiment EPSP sizes at baseline and final 10 minutes (Wilcoxon signed ranks: NRe Z = −2.7, p = 0.008; HPC Z = −2.7, p = 0.008). Normalised final EPSP amplitudes for −10 ms delay pairing performed at −70 and −50 mV (2-way ANOVA, effect of membrane potential: F_(1,28)_ = 11.3, p =0.0023, main effect of pathway: F_(1,28)_ = 0.005, p = 0.95, interaction: F_(1,28)_ = 0.14, p = 0.7). N = 9 cells from 9 animals **D** – Pairing with +10 ms lag (NRe precedes HPC by 10 ms) with depolarisation does not induce plasticity in either pathway (Wilcoxon signed ranks NRe: Z = −0.14, p = 0.89, HPC: Z = −0.42, p = 0.67). Traces as in **B**. Normalised final EPSP amplitudes for +10 ms and −10 ms at −50 mV membrane potential (2-way ANOVA, main effect of pairing order: F_(1,30)_ = 5.2, p = 0.030, main effect of pathway: F_(1,30)_ = 0.12, p = 0.73, interaction: F_(1,30)_ = 0.0002, p = 0.99). N = 8 cells from 7 animals. **E** – Bath application of NMDA receptor antagonist D-AP5 (50 µM), as indicated by grey shading, blocks induction of LTD by −10 ms pairing. Traces as in **B**, except EPSP scale bars = 1 mV. (Wilcoxon signed ranks NRe: Z = −0.84, p = 0.4; HPC Z = −1.3, p = 0.21). Normalised amplitudes for −50 mV pairing in absence or presence of D-AP5 (2-way ANOVA, main effect of drug: F_(1,.30)_=21.0, p < 0.0001, main effect of pathway: F_(1,30)_ = 0.9, p = 0.35, interaction: F_(1,30)_ = 2.6, p = 0.12). N = 8 cells from 8 animals.

We performed pairing of NRe and HPC (10 ms ISI) repeated (100 pairs) at 5 Hz with the cell held at −70 mV membrane potential. Pairing consisted of either HPC preceding NRe (−10 ms lag), or NRe preceding HPC (+10 ms lag). Under these conditions pairing was exclusively subthreshold in all cells and did not result in plasticity when HPC led NRe (−10 ms) or vice versa (+10 ms; Fig 7B, S6A). The same results were obtained when ISI was reduced to ± 5ms (Fig S6A). As NRe has also been proposed to be involved in coordinating slow oscillations in the 0.1-1 Hz band (Dolleman-van der Weel et al 2019) and ITDP delivered at 1Hz induces plasticity in hippocampus (Dudman et al 2007), we repeated ITDP protocols at 1Hz, however these again failed to induce synaptic plasticity regardless of the pairing windows used (Fig S6A).

To introduce spiking to the postsynaptic cell and increase NMDA receptor activation we modified the 5 Hz pairing protocol by depolarising cells to −50 mV by intracellular current injection for the duration of the pairing. Under these conditions when HPC fibres were stimulated 10 ms before NRe (−10 ms lag) pairing produced LTD in both NRe (p = 0.008) and HPC (p = 0.008) inputs (Fig 7C) and was significantly different to the same protocol applied at −70 mV two-way ANOVA of pairing at −70 and −50 mV: main effect of membrane potential p = 0.0023). To account for the emergence of LTD when pairing was delivered at −50 mV, we examined cell firing rates during pairing. At −50 mV, pairing produced a modest number of spikes which occurred as single as opposed to burst events (Fig S6D; 21.9 ± 7.5 spikes from 100 pairings, compared to none at −70 mV) which may account for the presence of plasticity.

Next, we tested the hypothesis that plasticity depends on the temporal order of synaptic inputs by reversing the order of stimulation such that NRe preceded HPC by 10 ms (+10 ms). Changing the order of pairing did not affect the number of spikes (Fig S6D), but surprisingly this protocol did not induce plasticity in either input (Fig 7D; NRe p = 0.89, HPC p = 0.67, two-way ANOVA of +10 vs −10 ms: main effect of pairing order: p = 0.030). These data show that associative plasticity of NRe and HPC synaptic inputs into mPFC critically depends on the temporal order of the incoming afferents.

Having shown that unidirectional pairing induces LTD in depolarised cells, each pathway was then stimulated alone at −50 mV. Stimulation of either HPC (Fig S6B; p = 0.23) or NRe fibres alone (Fig S6C; p = 0.40) did not induce plasticity in the test or control pathway, confirming that this form of plasticity is associative in nature. Spiking was not significantly different to that with the pairing protocol applied in either of these experiments (Fig S6D).

Associative plasticity of NRe and HPC is dependent upon depolarisation of the postsynaptic cell to −50 mV, resulting in spiking and presumably greater NMDA receptor activation. To identify the mechanisms by which plasticity is mediated we therefore paired HPC and NRe stimulation (−10 ms) at −50 mV in the presence of NMDA receptor antagonist D-AP5 (50 µM; Fig 7E). Neither NRe (p = 0.4) nor HPC (p = 0.21; two-way ANOVA of −10 ms pairing vs D-AP5 data, main effect of drug: p <0.0001) underwent plasticity in the absence of NMDAR activity. No significant difference in spiking was observed due to the presence of D-AP5 (Fig S6D), furthermore numbers of spikes evoked during pairing was not a good predictor of the degree of LTD in the −10 ms, −50 mV experiments (data not shown). Together these data suggest that elevated NMDA receptor activation rather than spiking per-se is a key determinant of plasticity. Together these results show that during depolarisation NRe and HPC inputs interact in a unidirectional manner via NMDAR-mediated transmission to induce an associative form of synaptic plasticity at layer V mPFC pyramidal neurons

## Discussion

Thalamic NRe has emerged as an additional important brain region for performance of higher order-cognitive tasks which require HPC-mPFC interactions. Here we advance the understanding of this circuit, showing that NRe and HPC inputs converge onto L5 pyramidal neurons in prelimbic cortex, these inputs undergo markedly different short-term plasticity and neuromodulation via muscarinic acetylcholine receptors, and interact with specific timing and directionality to induce associative synaptic plasticity, revealing a potential memory encoding mechanism.

Anatomical evidence shows NRe axon labelling across all layers of mPFC (Vertes et al 2006) and i*n vivo* field recordings have recorded large amplitude EPSPs in both superficial and deep layers of prelimbic cortex following NRe stimulation (Di Prisco & Vertes 2006, Eleore et al 2011). Our data show that a high proportion (75%) of L5 pyramidal neurons receive input from NRe and demonstrate with layer and cell-type specificity that NRe fibres synapse directly onto L2/3 and L5 pyramidal cells. These data advance current understanding of the tripartite HPC-NRe-mPFC circuitry.

Previous studies have coalesced electrophysiological and anatomical data to show that L5 pyramidal neurons with different projection targets have different intrinsic membrane properties, (Anastasiades et al 2018, Dembrow et al 2010, Dembrow et al 2015, Gee et al 2012). In the present study the parameters of cells receiving NRe input alone or NRe and HPC inputs most closely resembled intratelencephalic (IT) cells also recorded from L5 of PFC in rat (Dembrow et al 2010) having median input resistance >100 MΩ and sag <10 % (Fig 1E), although a minority of cells do express strong sag current suggesting their targets are not exclusively IT cells. Significant differences were found between input resistance and sag of cells which had only HPC input compared to those innervated by NRe. As HPC afferents have themselves previously been shown to preferentially target IT over pyramidal tract (PT) neurons in L5 (Dembrow et al 2015, Liu & Carter 2018) the fact that cells receiving NRe input more closely resemble IT neurons than those which only have HPC input suggests that NRe also preferentially targets IT cells. In this respect NRe is similar to mediodorsal (MD) thalamic input to mPFC which also preferentially targets IT neurons (Collins et al 2018). Further investigation is required to confirm cell-specific targeting by NRe. Overall, it is likely that convergent NRe and HPC input results in both local feed-forward excitation from L2 pyramidal projections to L5 pyramidal neurons (Collins et al 2018), inter-hemispheric cortico-cortical excitation via L2 and L5 IT cells and some degree of subcortical output via the lesser targeted PT cells. Our findings show that NMDA receptors at NRe inputs, like those of HPC inputs, have prominent GluN2B-subunit expression (Flores-Barrera et al 2014). The combination of the slow kinetics of GluN2B-containing NMDA receptor currents and targeting of principal neurons with intra- and cross-hemispheric projections position NRe and HPC inputs to readily recruit persistent firing patterns to support working memory (Hallock et al 2016, Monaco et al 2015).

### Short-term plasticity of NRe-mPFC transmission

A novel finding of the current study is that at theta-frequencies, which correspond to the instantaneous firing rate of NRe matrix cells (Walsh et al 2017), NRe input to both L2/3 and L5 pyramidal cells undergoes strong short-term depression (Fig 3A,B). This contrasts the facilitation reported in *in vivo* studies (Di Prisco & Vertes 2006, Eleore et al 2011). However, in those studies intra-thalamic stimulation was delivered, which may mean that NRe neurons, rather than the synapses to mPFC, are the locus of the reported facilitation. *In vitro* data from a study of matrix thalamus to mPFC L2/3 pyramidal neurons (Cruikshank et al 2012) reported weak facilitation at 10 Hz; however only 1 of 11 animals in the data set reported had NRe ChR2 expression, with the remainder largely restricted to VM and AM thalamus, which have previously been reported to facilitate (Collins et al 2018). Our NRe data more closely resemble short-term depression seen from the mediodorsal thalamus (Collins et al 2018). HPC inputs to mPFC meanwhile do not undergo notable short-term plasticity at theta frequency (Fig 3A), in keeping with previous findings (Liu & Carter 2018).

What mechanism underlies short-term depression of NRe inputs? As HPC ChETA_TC_ stimulation at 5 and 10 Hz replicated electrical stimulation of HPC afferents (Fig 3C), it is unlikely that NRe short-term depression is due to artefacts of optogenetic stimulation or viral transduction (Jackman et al 2014). Differential AMPAR or NMDAR expression does not explain differences in summation as measures of transmission via these receptors did not differ. Furthermore, influence of mGlu, GABA_B_, nACh or presynaptic NMDA receptors does not appear to underlie NRe short term dynamics as blocking these receptors had no effect on theta-frequency synaptic transmission. Use dependent blockade of NMDAR currents with MK801 suggest that the short-term depression seen in NRe inputs is due to a high probability of release, as seen in other thalamocortical synapses (Gil et al 1999). In this respect NRe projections are alike those of primary sensory thalamus and MD-mPFC projections (Collins et al 2018, Sherman 2016).

During spatial working memory, analysis of the directionality of HPC-mPFC theta coherence has been shown to change from HPC leading mPFC during the delay phase, to mPFC leading HPC during a decision phase (Hallock et al 2016). If NRe assists in entraining mPFC to HPC theta, short-term depression of NRe inputs to mPFC may assist in the transition between bottom-up and top-down processing whereby mPFC is known to send trajectory information back to HPC via NRe (Ito et al 2015).

### Neuromodulation of NRe and HPC inputs

Cholinergic modulation of NRe and HPC synapses onto mPFC cells was strikingly different. Previous data have shown cholinergic activation of mPFC with CCh produces depression of locally evoked excitatory transmission (Caruana et al 2011, Huang & Hsu 2010, Martin et al 2015) and of HPC synapses onto both L2/3 and L5 pyramidal cells (Ghoshal et al 2017, Maksymetz et al 2019, Wang & Yuan 2009). Here we find that lower concentrations of CCh than used in the above studies acting via M2 muscarinic receptors produced acute depression of HPC input to L5 pyramidal cells.

In contrast to HPC transmission, NRe inputs were unaffected by CCh. This is surprising since NRe itself is abundant in both muscarinic and nicotinic receptors (Clarke et al 1985, Wamsley et al 1984) and infusion of muscarinic or nicotinic receptor antagonists into NRe results in deficits of associative recognition memory encoding (Barker & Warburton 2018). Other corticothalamic synapses have been shown to be potentiated by addition of nAChR agonists (Gil et al 1997) and nicotinic agonists have been shown to increase spontaneous excitatory transmission in mPFC, an effect which is absent from animals with extensive thalamic lesions (Lambe et al 2003). It is plausible that the relatively slow bath application of CCh used in the present study does not capture the effects of rapidly desensitising nAChRs. Muscarinic-LTD is also absent in MD projections to mPFC L5 (Maksymetz et al 2019), suggesting that insensitivity to muscarinic LTD may be a feature of thalamic inputs to mPFC.

Previous reports have shown varied effects on glutamatergic transmission in mPFC, including both potentiation and depression of AMPAR and NMDAR mediated components, mediated via D1Rs and D2Rs (Banks et al 2015, Gao et al 2001, Gonzalez-Islas & Hablitz 2003, Law-Tho et al 1994, Seamans et al 2001, Urban et al 2002). In this study neither NRe or HPC AMPA EPSPs at −70 mV were affected by either D1R or D2R agonists. In similar experiments we have previously shown that HPC NMDARs undergo D2R-dependent depression (Banks et al 2015) but there was no direct effect on AMPAR transmission (though see: Burke et al 2018). Dopamine’s role in the mPFC may lie in its effects on local synaptic transmission (Burke et al 2018), cell excitability (Anastasiades et al 2019, Gee et al 2012), and modulation of synaptic plasticity (Otani et al 2003).

### Input-timing dependent plasticity of NRe and HPC synapses

Here we report a novel form of associative plasticity at both the HPC and NRe synapses induced by pairing of these inputs, but only when HPC leads NRe. The plasticity protocol was designed to be physiologically plausible, with the timing windows predicted on a simplified version of the HPC-NRe-mPFC tripartite circuit (Dolleman-van der Weel et al 2019). Synaptic plasticity was not induced at resting membrane potential but was induced at −50 mV and was blocked by D-AP5, suggesting that NRe and HPC synapses interact via NMDARs to induce plasticity. In cortex, NMDA receptors are tetramers composed of two obligatory GluN1 subunits and any permutation of two GluN2A/GluN2B subunits. GluN2B containing receptors, which have slower kinetics than those only expressing GluN2A subunits (Paoletti et al 2013), are expressed at higher levels in adult mPFC compared to other cortical regions (Wang et al 2008) including at HPC afferents (Flores-Barrera et al 2014). Our findings show that NRe and HPC synapses had equivalent levels of NMDAR expression and similar sensitivity to a GluN2B-selective antagonist, suggesting that both these inputs are abundant in GluN2B subunits, which is not a universal feature of synapses onto L5 pyramidal neurons (Flores-Barrera et al 2014). GluN2B subunit kinetics facilitate sustained charge-transfer, and therefore Ca^2+^ influx, at low frequency (Erreger et al 2005) which contributes to GluN2B-dependent LTD (Massey et al 2004). Slow NMDAR activity may be conducive to interaction of the spines of NRe and HPC synapses via spatiotemporal summation with possible mechanisms for plasticity induction including activation of calcium-dependent second messengers, release of calcium from intracellular stores (Dudman et al 2007) or generation of dendritic calcium spikes (Larkum 2013). It remains unclear as to whether generation of action potentials is required for input-timing dependent plasticity (ITDP): although spiking was elicited by −10 ms pairing at −50 mV the number of spikes did not correlate with the degree of plasticity, nor was there any difference in number of spikes when pairing order was reversed. Together, this suggests that action potentials are not a critical factor in induction of ITDP, as has also been described for ITDP in other brain regions (Dudman et al 2007, Williams & Holtmaat 2019).

ITDP of NRe and HPC inputs is, to the best of our knowledge, distinctive in that plasticity was induced in both pathways, in contrast to that in other brain regions where only the more proximal of the two synaptic inputs undergoes plasticity (Dudman et al 2007, Williams & Holtmaat 2019). This suggests that interaction of HPC and NRe synapses during pairing occurs in overlapping dendritic components, thus promoting spread of NMDAR-mediated depolarisation and equalising Ca^2+^ influx between spines. Functional and anatomical data supports overlapping distribution of synaptic input from HPC (Liu & Carter 2018) and NRe (Fig 1B) in deep layers of mPFC. In addition, NRe and HPC EPSPs show equal rise time (NRe 3.6 ± 0.3 ms, HPC 3.7 ± 0.3 ms) indicating equal distance from the soma (Sjostrom & Hausser 2006).

However, the mechanism by which pairing induced plasticity when HPC stimulation preceded NRe but not when pairing order was reversed are not clear. Possible explanation may involve differential activation of feed-forward inhibitory circuits, for example NRe may recruit stronger feedforward inhibition than HPC which may impair summation between pathways when NRe activation precedes HPC. Alternatively feedforward inhibition recruited by NRe and HPC could target different subcellular locations to achieve the same effect (Cruikshank et al 2012) or activation of HPC inputs could result in tightly timed disinhibition to allow plasticity with one direction of pairing but not the other (Williams & Holtmaat 2019). To fully understand which of these (combinations of) possibilities is important will require considerable further investigation.

Each of mPFC, HPC and NRe are required for many high order mnemonic and executive functions (Dolleman-van der Weel et al 2019) including spatial navigation (Ito et al 2015, Jankowski et al 2015), associative recognition memory (Barker & Warburton 2018) and sequence memory (Jayachandran et al 2019), but the circuit mechanisms underlying these functions are not understood. Whilst the behavioural function of the associative plasticity we describe remains to be determined, the specific timing conditions for plasticity we report suggest that NRe may impose a timing control which determines the salience of HPC signals and promotes their encoding. NRe has been noted to receive input from many other higher order regions including but not limited to entorhinal cortex, perirhinal cortex and amygdala (Dolleman-van der Weel et al 2019), and could act to integrate, for example, novelty or contextual information with incoming HPC spatial information which may promote encoding or consolidation of object-place associations (Barker & Warburton 2018). Such a mechanism may depend on NRe to coordinate oscillatory activity across multiple regions, and the observation of a high incidence of connectivity between NRe and L5 principal neurons in the present study may act to amplify signals from perirhinal and entorhinal cortices which have more sparse direct connections with prelimbic cortex (Hoover & Vertes 2007) but which are crucial for many forms of memory processing.

## Acknowledgments

This work was funded by Wellcome Trust Grant 206401/Z/17/Z and BBSRC grant BB/L001896/1. We would like to thank Jack Mellor, Paul Anastasiades and members of the Bashir and Warburton laboratories for contributions, helpful discussion and comments on the manuscript, and Dr Clair Booth for assistance with MATLAB analyses and mCherry staining.

## Notes

### Competing Interest Statement

The authors have declared no competing interest.

