## Supplementary figures for "Plasticity in prefrontal cortex induced by coordinated nucleus reuniens and hippocampal synaptic transmission"

**Supplementary Figure 1**

**A** – Action potential properties measured from first spike fired at the lowest current injection level which produced spiking in Fig S1B. Threshold: ANOVA $F_{(3,183)}$= 4.0, p = 0.0085, * p < 0.05 Tukey's multiple comparisons. Other parameters tested using Kruskal-Wallis due to one or more column failing Shapiro-Wilk test for normality. Peak, p = 0.66. Max rate of rise, p = 0.069. Width, p = 0.19.

**B –** Left panel, number of spikes fired in response to 500 ms depolarising current injections of varying amplitudes, 2-way ANOVA main effect of synaptic input $F_{(3,183)}$ = 0.43, p = 0.74. Right, instantaneous firing frequency for first 7 action potential pairs, repeated-measures ANOVA between subjects effect of synaptic input $F_{(3,182)}$ = 1.2, p = 0.30. Data shown as mean ± SD, n as indicated in Fig 1E.

**C** – Left panel, medium afterhyperpolarisation (mAHP) amplitude following fixed numbers of action potentials. Between subject effects: $F_{(3,182)}$ = 1.0, p = 0.38. Centre: afterdepolarisation (ADP) amplitude following single action potentials evoked by 2000 pA, 2 ms current. Kruskal-Wallis p = 0.085. Right, percentage of cells in each group with prevalent ADP, Pearson $\chi^2$ = 4.95, p = 0.18.

**D** – Basal NRe and HPC EPSPs are blocked by bath application of tetrodotoxin (0.5 µM). Increasing stimulus strength or application of 4-AP (100 µM) alone were insufficient to rescue NRe transmission, however increased stimulation strength in the presence of 4-AP restored NRe transmission. These data are consistent with basal optogenetic stimulation parameters producing action-potential dependent EPSPs, with increased stimulus strength required to achieve over-bouton release in 4-AP. NRe EPSPs were completely blocked by glutamate antagonists. Note that electrically evoked HPC EPSCs are not inducible even in the presence of 4-AP. NRe stimulus strength was achieved by increasing light pulse duration to 5ms from baseline values of (0.2-2ms). HPC stimuli were doubled in amplitude compared to baseline.

**E** – EPSP traces from an example experiment in (**D**).

**F** – HPC EPSC blocked by bath application of 5 µM NBQX.

**Supplementary Figure 2 – Optogenetic stimulation for high-frequency synaptic transmission**

**A** Short-term depression of NRe EPSPs at 5 and 10 Hz shows no relationship to duration of light pulses used to excite $ChETA_{TC}$-expressing NRe afferents. Blue line shows linear regression, dotted lines 95% confidence intervals (slope: 5 Hz = -0.054 ± 0.11, 10 Hz = -0.069 ± 0.12). Data taken from baselines shown in Fig 3A, 4B, and S3A-D, n = 42.

**B** Comparison of HPC afferent electrical and optogenetic stimulation. Top row: mean ± SEM of normalised amplitudes showing that $ChETA_{TC}$ was unable to reproduce the short-term plasticity of electrical stimulation at 20 Hz (main effect of stimulation method 20 Hz: $F_{(1,12)} = 10.3$, $p = 0.008$; response number: $F_{(2.2,26.9)} = 14.8$, $p = 3 \times 10^{-5}$; interaction: $F_{(2.7,32.2)} = 3.2$, $p = 0.041$), 50Hz (stimulation method $F_{(1,12)} = 9.45$, $p = 0.01$; response number $F_{(1.6,19.2)} = 21.2$, $p = 3 \times 10^{-5}$; interaction $F_{(2.1,24.7)} = 4.8$, $p = 0.016$) or 100 Hz stimulation (stimulation method $F_{(1,12)} = 30.2$, $p = 0.0003$; response number $F_{(1.4,14.1)} = 12.1$, $p = 0.002$; interaction $F_{(1.7,17.3)} = 6.3$, $p = 0.011$; n = 13). Bottom row: examples traces of optogenetic (green) and electrical (black traces) from same cell. Note lack of subsequent peaks after the first optogenetic response at 100 Hz, accentuated by the pronounced LED-artefacts in this recording. Blue squares indicate LED activation. Scale bars = 3 mV/50 ms except 20 Hz, where x-axis is 100 ms.

**Supplementary Figure 3 – Antagonism of group II mGlu, $GABA_B$, NMDA or nicotinic receptors does not affect short-term plasticity of NRe or HPC synapses at theta frequencies**

**A** Bath application of group II mGluR receptor antagonist EGLU (10 µM) did not affect transmission in either pathway at 5 or 10 Hz (5 Hz: NRe $F_{(8,72)} = 0.48$, p = 0.87; HPC $F_{(3.1, 28)} = 0.98$, p = 0.42; 10 Hz: NRe $F_{(3.3,29.9)} = 2.8$, p = 0.053, HPC $F_{(3.3, 29)} = 2.55$, p = 0.071).

**B** $GABA_B$ receptor antagonist CGP55845 (1 µM) did not affect short-term plasticity of NRe or HPC at 5 or 10Hz (5 Hz: NRe $F_{(8,72)} = 1.3$, p = 0.28; HPC $F_{(1.9,17.3)} = 0.51$, p = 0.60; 10 Hz: NRe $F_{(8,72)} = 0.5$, p = 0.86, HPC $F_{(2.9, 26.2)} = 0.80$, p = 0.50).

**C** NMDAR antagonist D-AP5 (50 µM) did not alter short-term plasticity at 5 or 10 Hz in either pathway (5 Hz: NRe $F_{(1.9,11.2)} = 1.1$, p = 0.38; HPC $F_{(1.9,9.6)} = 0.45$, p = 0.64; 10 Hz: NRe $F_{(1.5,8.9)} = 2.5$, p = 0.15, HPC $F_{(2.0, 10.1)} = 1.8$, p = 0.22).

**D** Bath application of nicotinic acetylcholine receptor antagonist mecamylamine (1 µM) did not affect transmission in either pathway at 5 or 10 Hz (5 Hz: NRe $F_{(2.5,20.3)} = 1.42$, p = 0.27; HPC $F_{(7.6, 60.8)} = 0.82$, p = 0.52; 10 Hz: NRe $F_{(3.1,24.4)} = 0.35$, p = 0.79, HPC $F_{(2.3, 18.7)} = 0.9$, p = 0.44).

For all statistical tests Greenhouse-Geisser corrections were applied when Maulchly's test of sphericity was below 0.05

**Supplementary Figure 4 – Summary of cholinergic modulation of inputs to PFC**

**A**– Summary of CCh effects on normalised NRe and HPC EPSPs at acute (10-19 mins) and washout (40-49 mins) timepoints. Bars = mean +SEM, circles = individual cells. Two-way repeated-measures ANOVA revealed a timepoint*pathway*concentration interaction, demonstrating that CCh selectively attenuates HPC, not NRe inputs, in a reversible, concentration-dependent manner (main effects: pathway $F_{(1,20)} = 0.5$, $p = 0.48$; time-point $F_{(1.8,36)} = 11.3$, $p = 0.001$; concentration $F_{(2,20)} = 1.0$, $p = 0.37$; interactions: pathway x concentration $F_{(2,20)} = 1.2$, $p = 0.31$; timepoint x concentration $F_{(3.6,36)} = 3.9$, $p = 0.011$; pathway x timepoint $F_{(1.8,35)} = 14.7$, $p = 0.00004$; timepoint x pathway x concentration $F_{(4,40)}=3.0$, $p = 0.029$). Post-hoc analysis with Sidak's multiple comparison: **p = 0.0016; ***p = 0.0002. All other comparisons p > 0.05.

**B** – Summary of data shown in Fig4B & C. Two-way repeated-measures ANOVA (NRe: main effect of drug $F_{(2,34)} = 2.9$, $p = 0.07$; timepoint: $F_{(1,34)} = 0.0009$, $p = 0.98$; interaction $F_{(2,34)} = 0.18$, $p = 0.84$; HPC: main effect of drug $F_{(2,34)} = 9.5$, $p = 0.0005$; timepoint: $F_{(1,34)} = 12.8$, $p = 0.0011$; interaction $F_{(2,34)} = 1.3$, $p = 0.29$. Sidak's post-hoc analysis, *** p = 0.001. All other comparisons p > 0.05.

**Supplementary Figure 5 – Dopaminergic modulation of inputs to PFC**

**A -** Summary of SKF81297 data showing individual experiments normalised to baseline overlaid over mean ± SEM bar graph. * denotes significance vs baseline, $p = 0.0072$. Acute/washout are averaged amplitudes from final 10 mins of drug application and final 10 mins of recording.

**B** – Paired-pulse ratio (100 ms inter-stimulus-interval) is not altered by bath application of SKF81297 (10 µM). Two-way repeated-measures ANOVA: main effect of timepoint: $F_{(2,36)} = 0.1$, $p = 0.9$; pathway $F_{1,18)} = 0.44$, $p = 0.51$; interaction $F_{(2,36)} = 1.0$, $p = 0.37$. Left shows mean + SEM values, right shows individual experiment PPR in each pathway at baseline, final 10 minutes of drug application and final ten minutes of recording.

**C** – EPSP amplitudes in response to quinpirole for individual cells as shown in Fig 5C. NRe and HPC pathways shown in blue/black, respectively.

**D** – Acute effect of SKF81297 (10 µM) on synaptic strength plotted versus sum of sag and rebound in response to a -100 pA hyperpolarising current injection or cell input resistance at start of recording. Slopes are not significantly different from 0 in any instance. NRe and HPC pathways shown in blue/black, respectively.

**E** – Quinpirole (10 µM) data as for **D**. Slopes are not significantly different from 0 in any instance. NRe and HPC pathways shown in blue/black, respectively.

**Supplementary Figure 6 – Input-timing dependent plasticity of NRe and HPC does not occur at -70 mV and requires activation of both pathways**

**A** – Summary of pairing experiments showing mean + SEM normalised amplitude of NRe and HPC EPSPs 30-40 mins after pairing with different lags as indicated. Left graph shows pairing delivered at 5 Hz, right at 1 Hz. All experiments were performed at -70 mV. Plasticity was absent in both pathways at all frequencies and pairing delays tested (paired t-test of raw EPSP amplitudes, p values displayed on graph in blue for NRe and black for HPC). For all experiments n = number of cells, 1 cell per animal.

**B** – 5 Hz stimulation of HPC afferents at -50 mV (single stimuli) does not induce plasticity of test pathway (paired t-test, HPC $t_{(6)}$ = 1.4, p = 0.23) or control pathway (NRe, $t_{(6)}$ = 0.4, p = 0.74). Traces show example averaged EPSPs at baseline (blue/black) and 30-40 mins (red) and 15 pairings (grey), stimulation denoted by triangle. Scale bars EPSPs: 5 mV, 100 ms, pairing: 10 mV, 50 ms. Right panel: raw EPSP amplitudes at baseline and final 10 mins. N = 7 cells from 7 animals.

**C** – 5 Hz stimulation of NRe afferents at -50 mV (single stimuli) does not induce plasticity of test pathway (Wilcoxon signed ranks, NRe Z = -0.84, p = 0.40) or control pathway (HPC, Z = 0.98, p = 0.33). Traces show example averaged EPSPs at baseline (blue/black) and 30-40 mins (red) and 15 pairings (grey), stimulation denoted by triangle. Scale bars EPSPs: 2 mV, 100 ms, pairing: 10 mV, 50 ms. Right panel: raw EPSP amplitudes at baseline and final 10 mins. N = 7 cells from 7 animals.

**D** – Total number of spikes fired for experiments shown in Fig 7 and Fig S6. Data point represent individual cells, box plots show median, inter-quartile range and whiskers show minima and maxima. Data were subject to Kruskal-Wallis test (KW statistic = 14.6, p = 0.012) and post-hoc Dunn's multiple comparisons were carried out against vs -10 ms pairing at -50 mV, *** = 0.007, all other comparisons p > 0.05.

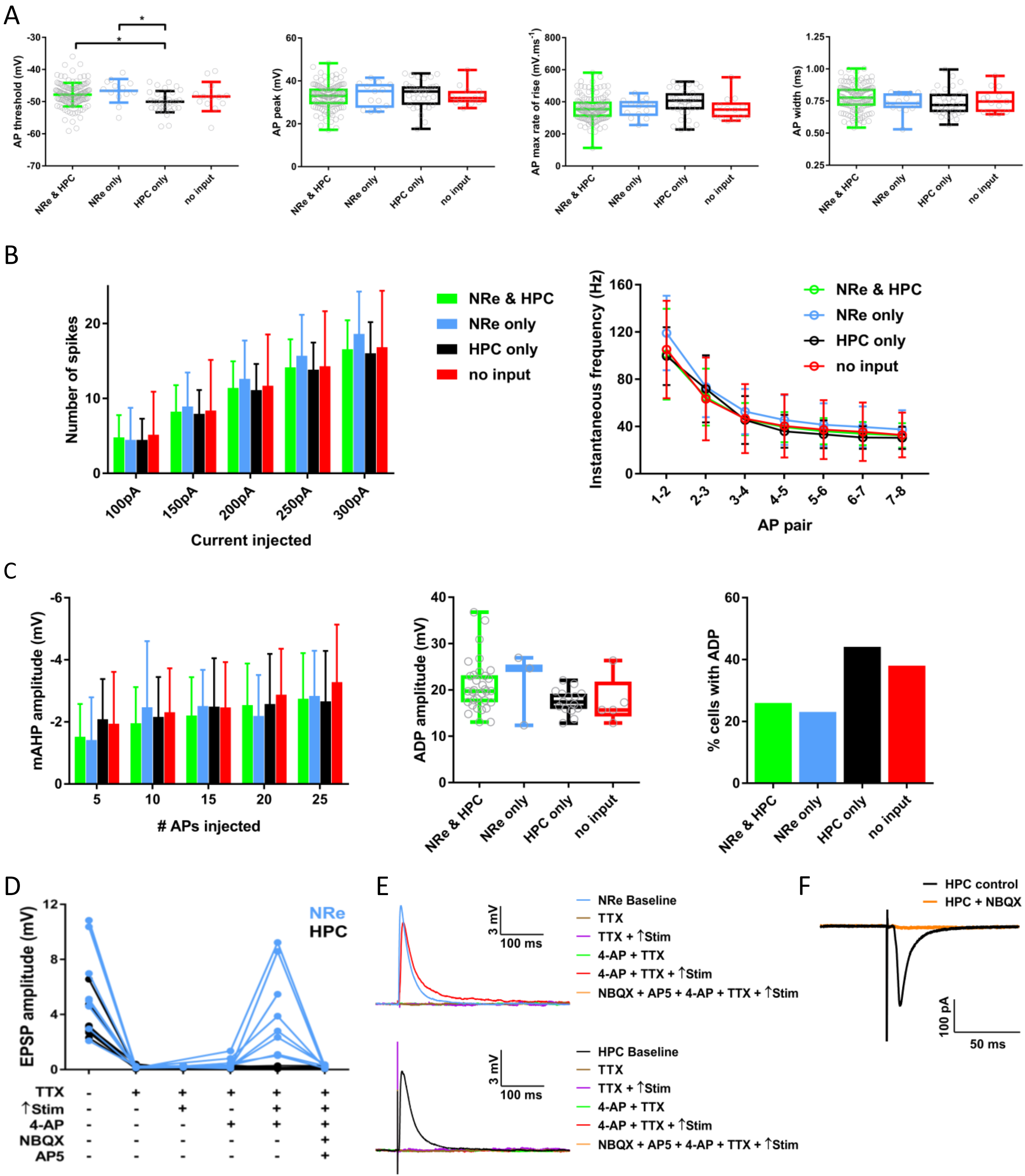

A
AP threshold (mV)
AP peak (mV)
AP max rate of rise (mV.ms-1)
AP width (ms)
NRe & HPC
NRe only
HPC only
no input
B
Number of spikes
Current injected
100pA
150pA
200pA
250pA
300pA
Instantaneous frequency (Hz)
AP pair
C
mAHP amplitude (mV)
### APs injected
ADP amplitude (mV)
% cells with ADP
D
EPSP amplitude (mV)
NRe
HPC
TTX
↑Stim
4-AP
NBQX
AP5
E
NRe Baseline
TTX
TTX + ↑Stim
4-AP + TTX
4-AP + TTX + ↑Stim
NBQX + AP5 + 4-AP + TTX + ↑Stim
HPC Baseline
3 mV
100 ms
F
HPC control
HPC + NBQX
100 pA
50 ms

A

B

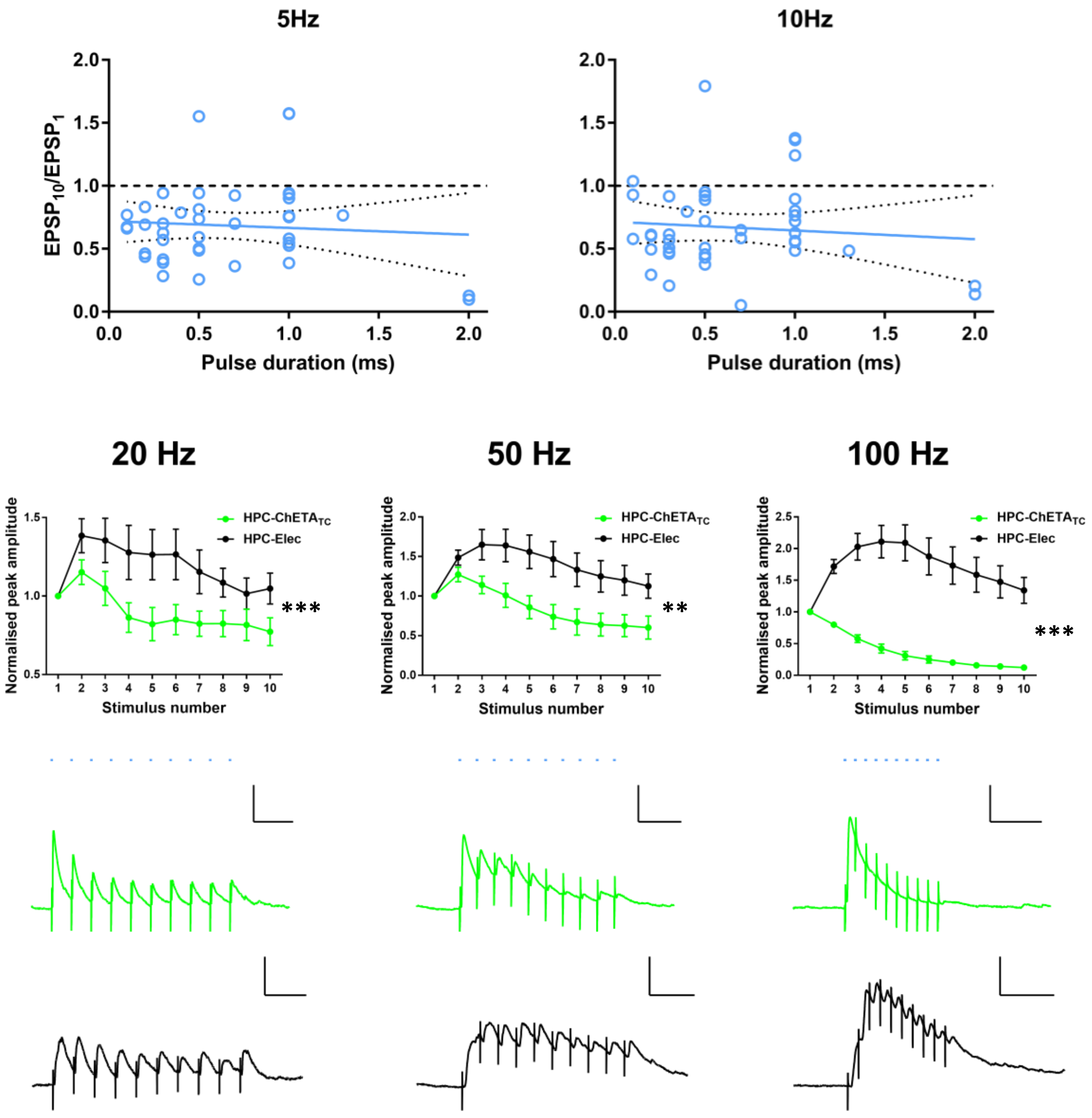
5Hz
10Hz
EPSP10/EPSP1
2.0
1.5
1.0
0.5
0.0
Pulse duration (ms)
20 Hz
50 Hz
100 Hz
Normalised peak amplitude
HPC-ChETATC
HPC-Elec
Stimulus number
***
**
***

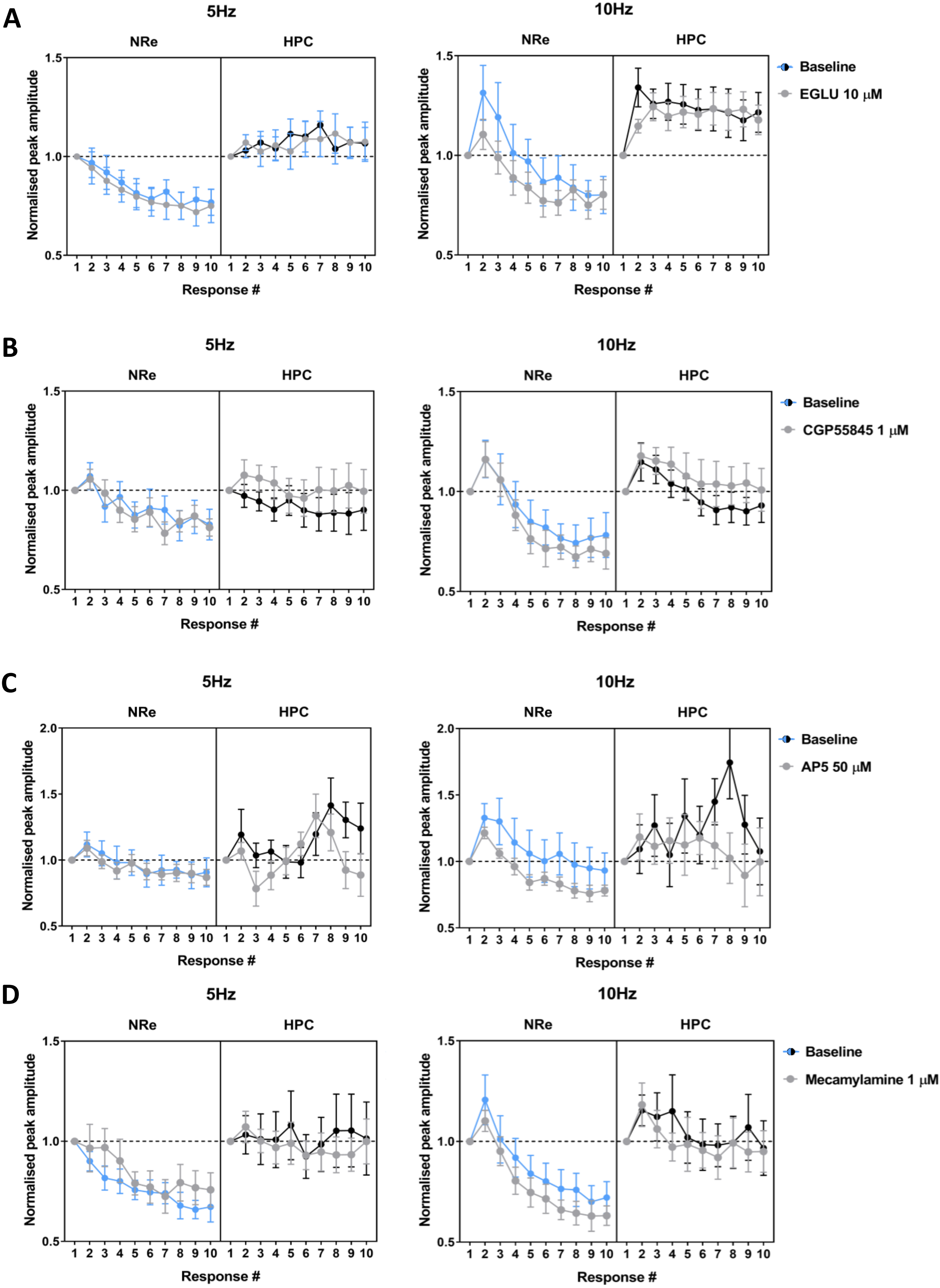

A
5Hz
10Hz
NRe
HPC
Normalised peak amplitude
Response #
Baseline
EGLU 10 μM
B
5Hz
10Hz
NRe
HPC
Normalised peak amplitude
Response #
Baseline
CGP55845 1 μM
C
5Hz
10Hz
NRe
HPC
Normalised peak amplitude
Response #
Baseline
AP5 50 μM
D
5Hz
10Hz
NRe
HPC
Normalised peak amplitude
Response #
Baseline
Mecamylamine 1 μM

A

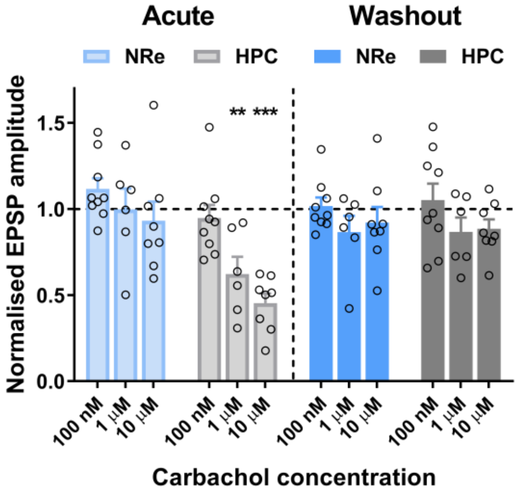

Acute
Washout
NRe
HPC
NRe
HPC
Normalised EPSP amplitude
0.0
0.5
1.0
1.5
** ***
100 nM
1 μM
10 μM
Carbachol concentration


B

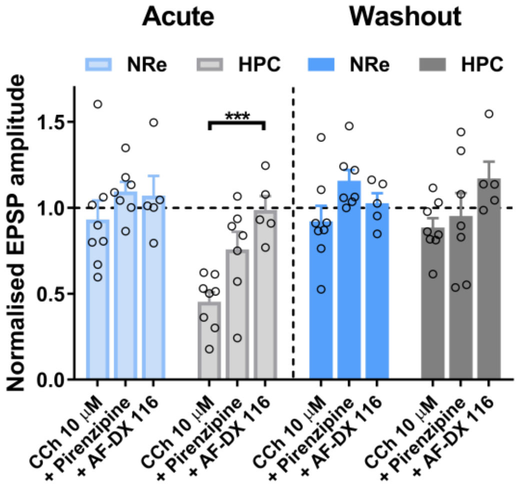

Acute
Washout
NRe
HPC
NRe
HPC
Normalised EPSP amplitude
0.0
0.5
1.0
1.5
***
CCh 10 μM
+ Pirenzipine
+ AF-DX 116

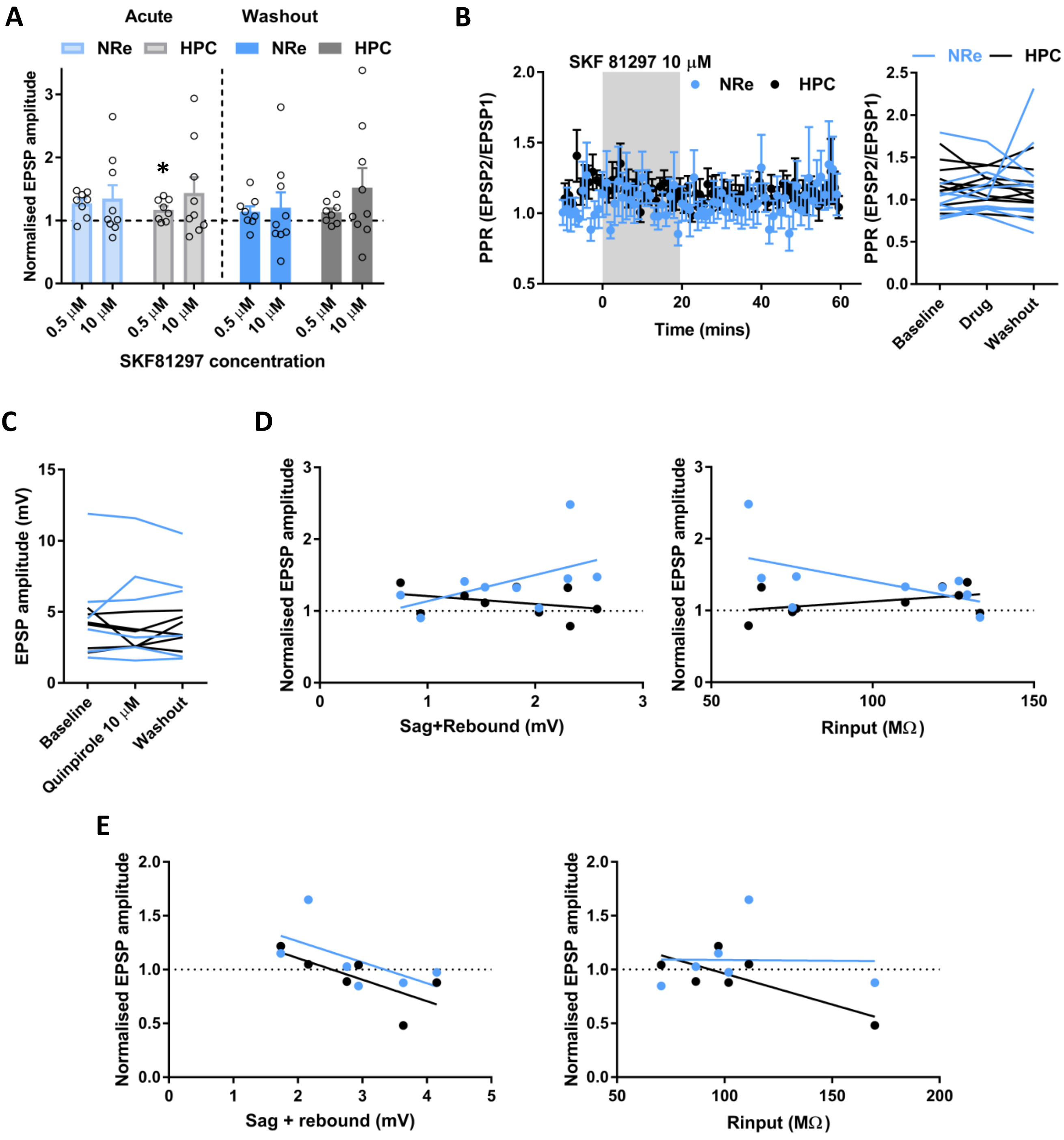
A
Acute
Washout
NRe
HPC
NRe
HPC
Normalised EPSP amplitude
0.5 μM
10 μM
SKF81297 concentration
B
SKF 81297 10 μM
NRe
HPC
PPR (EPSP2/EPSP1)
Time (mins)
Baseline
Drug
Washout
C
EPSP amplitude (mV)
Baseline
Quinpirole 10 μM
Washout
D
Normalised EPSP amplitude
Sag+Rebound (mV)
Rinput (MΩ)
E
Normalised EPSP amplitude
Sag + rebound (mV)
Rinput (MΩ)

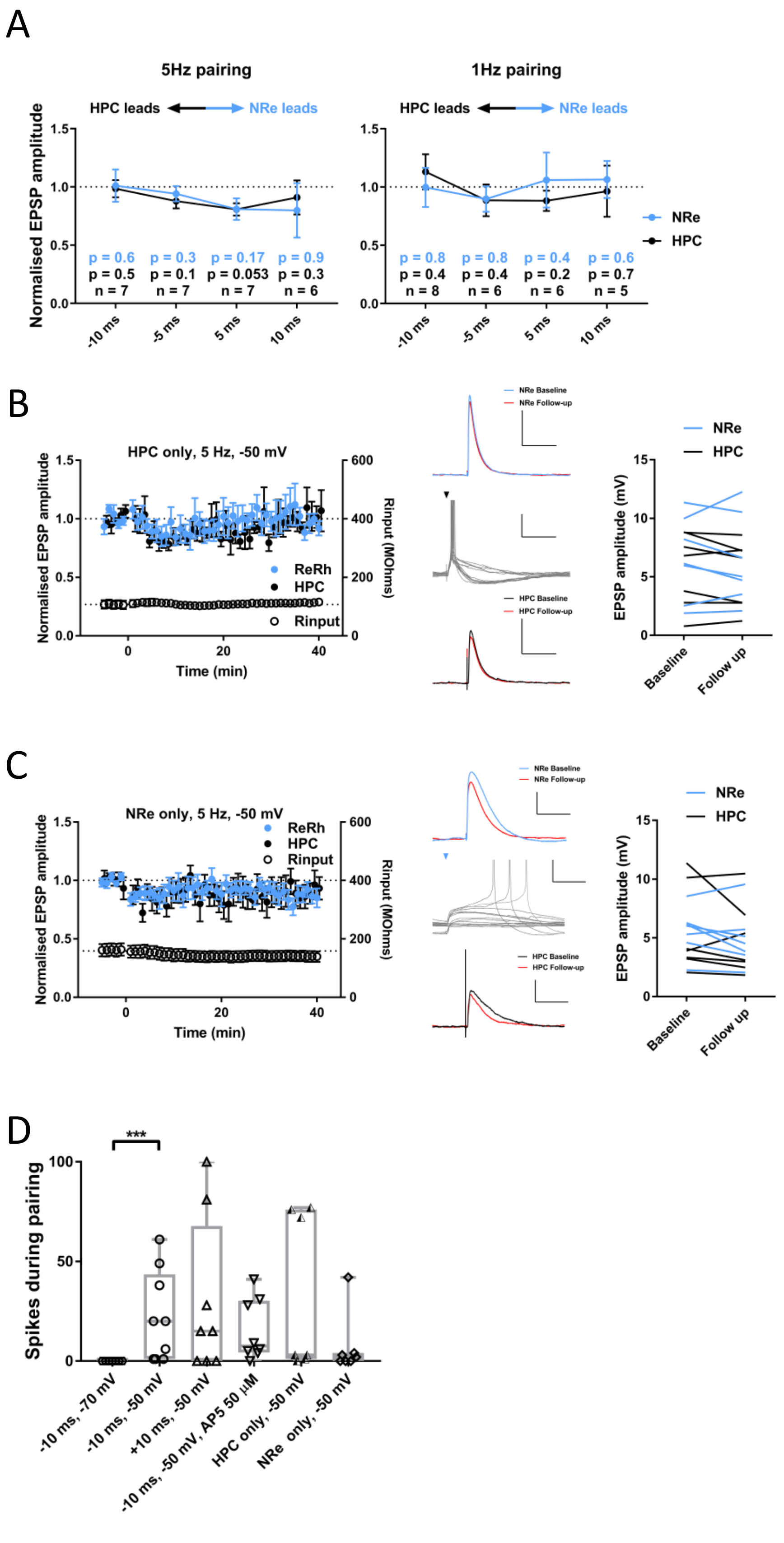

A
5Hz pairing
1Hz pairing
HPC leads
NRe leads
Normalised EPSP amplitude
p = 0.6
p = 0.5
n = 7
p = 0.3
p = 0.1
n = 7
p = 0.17
p = 0.053
n = 7
p = 0.9
p = 0.3
n = 6
p = 0.8
p = 0.4
n = 8
p = 0.8
p = 0.4
n = 6
p = 0.4
p = 0.2
n = 6
p = 0.6
p = 0.7
n = 5
-10 ms
-5 ms
5 ms
10 ms
NRe
HPC
B
HPC only, 5 Hz, -50 mV
ReRh
HPC
Rinput
Rinput (MOhms)
Time (min)
NRe Baseline
NRe Follow-up
HPC Baseline
HPC Follow-up
EPSP amplitude (mV)
Baseline
Follow up
C
NRe only, 5 Hz, -50 mV
D
Spikes during pairing
***
-10 ms, -70 mV
-10 ms, -50 mV
+10 ms, -50 mV
-10 ms, -50 mV, AP5 50 µM
HPC only, -50 mV
NRe only, -50 mV
